# Mosquito saliva sialokinin-dependent enhancement of arbovirus infection through endothelial barrier leakage

**DOI:** 10.1101/2021.02.19.431961

**Authors:** Daniella A Lefteri, Steven R Bryden, Marieke Pingen, Sandra Terry, Emily F Beswick, Georgi Georgiev, Marleen Van der Laan, Valeria Mastrullo, Paola Campagnolo, Robert Waterhouse, Margus Varjak, Andres Merits, Rennos Fragkoudis, Stephen Griffin, Kave Shams, Emilie Pondeville, Clive S McKimmie

**Affiliations:** Virus Host Interaction Team, Leeds Institute of Medical Research, School of Medicine, Faculty of Medicine and Health, University of Leeds, UK; MRC-University of Glasgow Centre for Virus Research, Glasgow, Scotland, United Kingdom; Inflammatory Skin disease group, Leeds Institute of Rheumatic and Musculoskeletal Medicine, School of Medicine, Faculty of Medicine and Health, University of Leeds, UK; Leeds Institute of Medical Research, School of Medicine, Faculty of Medicine and Health, University of Leeds, UK; Faculty of Health & Medical Sciences, Section of Cardiovascular Sciences, University of Surrey Guildford, UK; Department of Ecology and Evolution, Swiss Institute of Bioinformatics, University of Lausanne, 1015, Lausanne, Switzerland; Institute of Technology, University of Tartu, 50411 Tartu, Estonia; University of Edinburgh

**Keywords:** mosquitoes, arbovirus, inflammation, endothelium, vascular biology, zika, vaccines, skin

## Abstract

Viruses transmitted by *Aedes* mosquitoes constitute an increasingly important global health burden. Defining common determinants of host susceptibility to this large group of heterogenous pathogens are key for informing the rational design of new pan-viral medicines. Infection of the vertebrate host with these viruses is enhanced by the presence of mosquito saliva, a complex mixture of salivary gland-derived factors and microbiota. We show that enhancement of infection by saliva was dependent on vascular function and was independent of most anti-saliva immune responses, including to salivary microbiota. Instead, the *Aedes* gene product sialokinin mediated enhancement of virus infection through a rapid reduction in endothelial barrier integrity. Sialokinin is unique within the insect world as having vertebrate-like tachykinin sequence and is absent from non-vector competent *Anopheles* mosquitoes, whose saliva was not pro-viral and did not induce similar vascular permeability. Therapeutic strategies targeting sialokinin have potential to limit disease severity following infection with *Aedes* mosquito-borne viruses.

## Introduction

Mosquito-borne viruses are an important cause of debilitating and sometimes lethal infections. The most significant vectors of these viruses are *Aedes* species mosquitoes, which transmit several clinically important viruses (arthropod-borne, arboviruses), including dengue (DENV), Zika (ZIKV) and chikungunya (CHIKV) viruses (Weaver et al., 2018; Bhatt et al., 2013). Recent emergence of arboviruses into new geographic areas has led to explosive outbreaks of disease for which there are no licenced medicines or vaccines. Arboviruses are a large (>85 human pathogens), genetically diverse group of viruses that cause a wide spectrum of distinct diseases in humans (Gould and Solomon, 2008; Guzman and Harris, 2015; Weaver et al., 2018). This heterogeneity, combined with our inability to accurately predict the nature and timing of future epidemics, makes developing and stockpiling virus-specific drugs and vaccines in a timely manner challenging. It is therefore important that we better understand common determinants of host susceptibility to infection and thereby inform the rational design of new pan-viral medicines.

Factors that predispose the host to more severe arbovirus disease are poorly understood, but are most likely due to a combination of viral and host factors. This includes the host response to mosquito saliva at the skin inoculation site (Pingen et al., 2017; Fong et al., 2018; Conway et al., 2014b; Ribeiro and Francischetti, 2003; Schneider and Higgs, 2008; Schmid et al., 2016). Transmission of virus from biting mosquito to vertebrate host is a feature common to all mosquito-borne virus infections. Infected mosquitoes transmit virus to the mammalian host as they probe the skin for blood and deposit saliva, where virus replicates before disseminating via lymph to the blood. The mammalian host response to mosquito biting and/or mosquito saliva significantly enhances infection with many medically-important genetically-distinct viruses, including *Flaviviruses* e.g. DENV (Cox et al., 2012; McCracken et al., 2014; Schmid et al., 2016; Conway et al., 2014b) and West Nile virus (WNV) (Moser et al., 2016; Schneider et al., 2006; Styer et al., 2011), *Alphaviruses* e.g. Semliki Forest virus (SFV) (Pingen et al., 2016) and CHIKV (Agarwal e t al., 2016) and *Bunyavirales* e.g. Rift Valley fever virus, Cache Valley Virus and Bunyamwera virus (Edwards et al., 1998; Le Coupanec et al., 2013; Pingen et al., 2016).

Enhancement of virus infection by saliva is apparent within hours, resulting in a higher quantity of virus in tissues, blood and reduced survival (Pingen et al., 2017; Styer et al., 2011). As such, mosquito saliva is a key, common aspect of mosquito-borne virus infection that significantly worsens outcomes. Importantly, the mechanistic basis by which mosquito saliva increases host susceptibility to infection is poorly defined, although inflammatory responses that elicit an influx of virus-permissive monocytic cells are required (Pingen et al., 2016). Despite evidence that some pro-inflammatory mosquito salivary factors can enhance ZIKV infection (Hastings et al., 2019; Uraki et al., 2019) the ability of most salivary factors to modulate vertebrate susceptibility to virus are still being defined (Jin et al., 2018; Sun et al., 2020). Mosquito saliva is made up of a complex cocktail of bacterial microbiota and salivary gland gene encoded products (Ribeiro et al., 2007; Mancini et al., 2018; Ribeiro et al., 2010). Saliva facilitates mosquito blood feeding by promoting vasodilation and inhibiting various haemostatic processes, the effect of which on virus infection remain mostly uncharacterised.

We show here that a reduction in blood vessel barrier function, induced by *Aedes* saliva directly on endothelial cells, is necessary for enabling saliva-enhancement of arbovirus infection. We found that *Aedes* mosquito saliva only enhanced virus infection when applied *in vivo* and that the factor involved was independent of salivary microbiota, sensitive to heat denaturation and was only present in female mosquito saliva. Crucially, we found that *Anopheles* species mosquito saliva lacked the ability to enhance *Aedes* or *Anopheles*-borne virus infection and that this correlated with an inability to induce rapid endothelia barrier leakage. We exploited this difference to dissect the mechanistic basis by which *Aedes* saliva enhances infection with virus and to identify the requirement for the *Aedes* salivary factor sialokinin (SK). SK induced vascular leakage in skin and was both necessary and sufficient for defining *Aedes* saliva enhancement of infection. SK is absent in *Anopheles* and has high homology to mammalian substance P. As such, we define a novel aspect of disease transmission that can inform the development of novel vaccines with pan-viral potential that target *Aedes* salivary factors.

## Results

### Mosquito saliva is sufficient to enhance virus infection and worsen clinical outcome

Mosquito bites involves repeated skin tissue trauma and the deposition of saliva as the mosquito probes for a blood meal. Host responses to mosquito biting enhance incoming virus infection, as does the injection of experimentally-derived homogenates of salivary gland tissue (Pingen et al., 2017; Conway et al., 2014a). To define the factors and properties in isolated saliva responsible for modulating host susceptibility to co-inoculated virus in mice, we obtained saliva by forced salivation from adult *Ae. aegypti* female mosquitoes (Fig 1A,B and sFig1A). Saliva was co-injected into skin with either the *Alphavirus* SFV or *Flavivirus* ZIKV and virus RNA quantities in the skin inoculation site quantified by RT-QPCR, and virus infectious units determined in the serum, at 24 hours post infection (hpi). Although the presence of just one mosquito’s salivation was sufficient to induce a ten-fold increase in SFV (Fig 1A), this was variable, likely as a consequence of salivation efficiency between mosquitoes (sFig1B). Variation was much less pronounced when mice were co-injected with saliva obtained from 5 mosquitoes (Fig1A,B). Repeated tissue piercing with our hyperfine needles in the absence of injection, to simulate probing by mosquitoes, did not result in any virus enhancement (Fig 1C), suggesting enhancement of infection was dependent on saliva. Importantly, infection with saliva also reduced survival to SFV to a similar extent as injection of same titre virus into a mosquito bite (Fig 1D).

**Figure 1.**
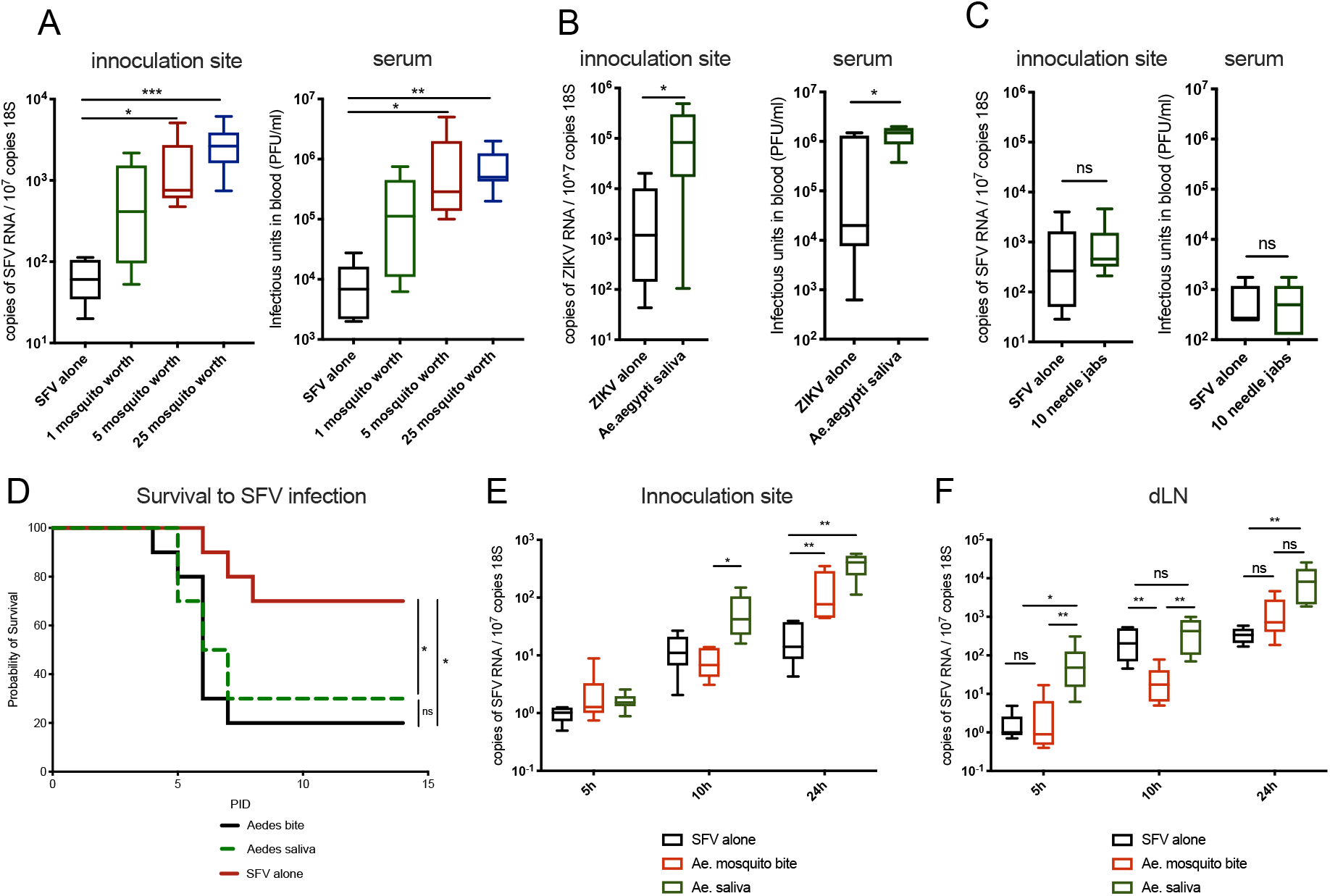
Mosquito saliva is sufficient to enhance virus infection and worsen clinical outcome. (A-F) Mice were inoculated with either 10^4^ PFU of SFV4 or 10^3^ PFU of ZIKV into skin of left foot (upper side), alone or following Ae.aegypti mosquito biting, or co-injected with Aedes saliva normalised for total protein content (mosquito salivated 0.3714μg protein on average). Viral RNA and host 18S were quantified by qPCR and viral titres of serum by plaque assays at 24hpi. (A) Mice were injected with SFV alone or alongside 1, 5 or 25 mosquitoes worth of saliva (n=6) (B) Mice were injected with ZIKV with or without 5 mosquitoes’ worth of saliva (1.86μg protein, n=6) (C) Mice were infected with SFV following one or 10 repeated tissue piercings with hyperfine needle. (n=6). (D) Survival of mice infected with 10^4^ PFU of SFV4 (n=10) (E and F) Mice were inoculated with SFV alone in resting skin, or into mosquito bitten skin (5 bites), or into resting skin mosquito saliva (1.86μg saliva protein, n=8)

Following infection of skin, some virus infects skin-resident cells, while some of the inoculum disseminates to the draining lymph node (dLN) where it also replicates (Pingen et al., 2016; Styer et al., 2011). Altered early dissemination of virus inoculum by saliva may explain enhancement of infection by saliva. However, while mosquito bites reduced dissemination of virus alone to dLN, mosquito saliva alone did not. This shows that while injection of virus into mosquito bitten skin, or co-injection of saliva with virus into resting skin, resulted in higher quantity of skin virus RNA, neither presence of bite or saliva alone enhanced early dissemination to dLN (Fig 1E, 1F).

Host inflammatory responses to mosquito biting that recruit virus-permissive monocytic cells are important for modulating host susceptibility to virus (Pingen et al., 2016). Here, we found that similar to mosquito biting, saliva alone in the absence of virus induced expression of pro-inflammatory genes *cxcl2*, *il1b* and *ccl2*, while prototypic anti-viral type I interferon (IFN) stimulated genes (*isg15* and *ccl5*) whose expression correlates with host resistance to infection (Bryden et al., 2020) were not altered (sFig 1C). In summary, we have optimised and defined our *in vivo* model for determining how saliva alone, in the absence of other mosquito bite or salivary gland tissue components, modulates host susceptibility to virus.

### Enhancement of virus infection *in vivo* requires processes that are absent *ex vivo*

To define whether mosquito saliva can directly modulate susceptibility of the two key cell types infected by SFV in the skin, we infected primary cultures of fibroblasts and macrophages *in vitro* with SFV in presence or absence of saliva. However, rather than enhancing infection, mosquito saliva either had no effect, or instead inhibited infection at some MOIs (Fig 2A and sFig 2A). This was a direct effect of saliva upon cells, rather than via direct action on the virus, as cells pre-treated with saliva for 20 minutes and washed prior to infection with SFV (referred to here as “saliva to cells”), also exhibited increased resistance to infection (sFig 2A and 2B). Interestingly, this was dependent on responses activated by bacterial microbiota, as microbiota-deficient saliva obtained from mosquitoes treated with broad-spectrum antibiotics (sFig 2C), was no longer able to mediate increased resistance to virus infection *in vitro* (Fig 2B). Antibiotics also rendered mosquito saliva less pro-inflammatory, as macrophages expressed significantly less of the prototypic pro-inflammatory chemokine *cxcl2* when exposed to treated saliva, in the absence of virus (Fig 2C). This lead us to hypothesise that activation of immune responses *in vitro* by microbiota may account for the ability of saliva to increase resistance to virus. Therefore, we decided to block signalling by the most important anti-viral pathway, type I IFN signalling, by the use of an IFN-receptor blocking antibody (Fig 2D). Here, incubation of cells with bacterial microbiota-containing saliva could no longer increase resistance to virus in IFN-R blocked cells. This demonstrated that, for *in vitro* cultured cells, salivary microorganisms could partially increase cellular resistance to virus in a type I IFN-dependent manner. Nonetheless, these *in vitro* studies failed to recapitulate the overall phenotype observed *in vivo,* in which virus infection was enhanced, suggesting the mechanism responsible overcomes localised bacterial innate responses at the injection site. In case cellular crosstalk by skin cells, or the presence of other cell types not present in skin explant cultures, were required to mimic the *in vivo* phenotype, we assessed the susceptibility of intact skin explants to infection, which we have previously used to study innate immune responses to arbovirus infection (Bryden et al., 2020). However, these explants did not exhibit any increase in susceptibility to virus following exposure to saliva. This included infection of explants derived from resting skin, mosquito bitten skin and explants derived from skin injected with saliva prior to biopsy (Fig 2E, 2F and sFig 2D). These cultured skin tissue or cells were not rendered more susceptible to infection by saliva, suggesting that a key *in vivo*-specific process, not present in these *ex vivo* systems, was required for the saliva to enhance virus infection.

**Figure 2:**
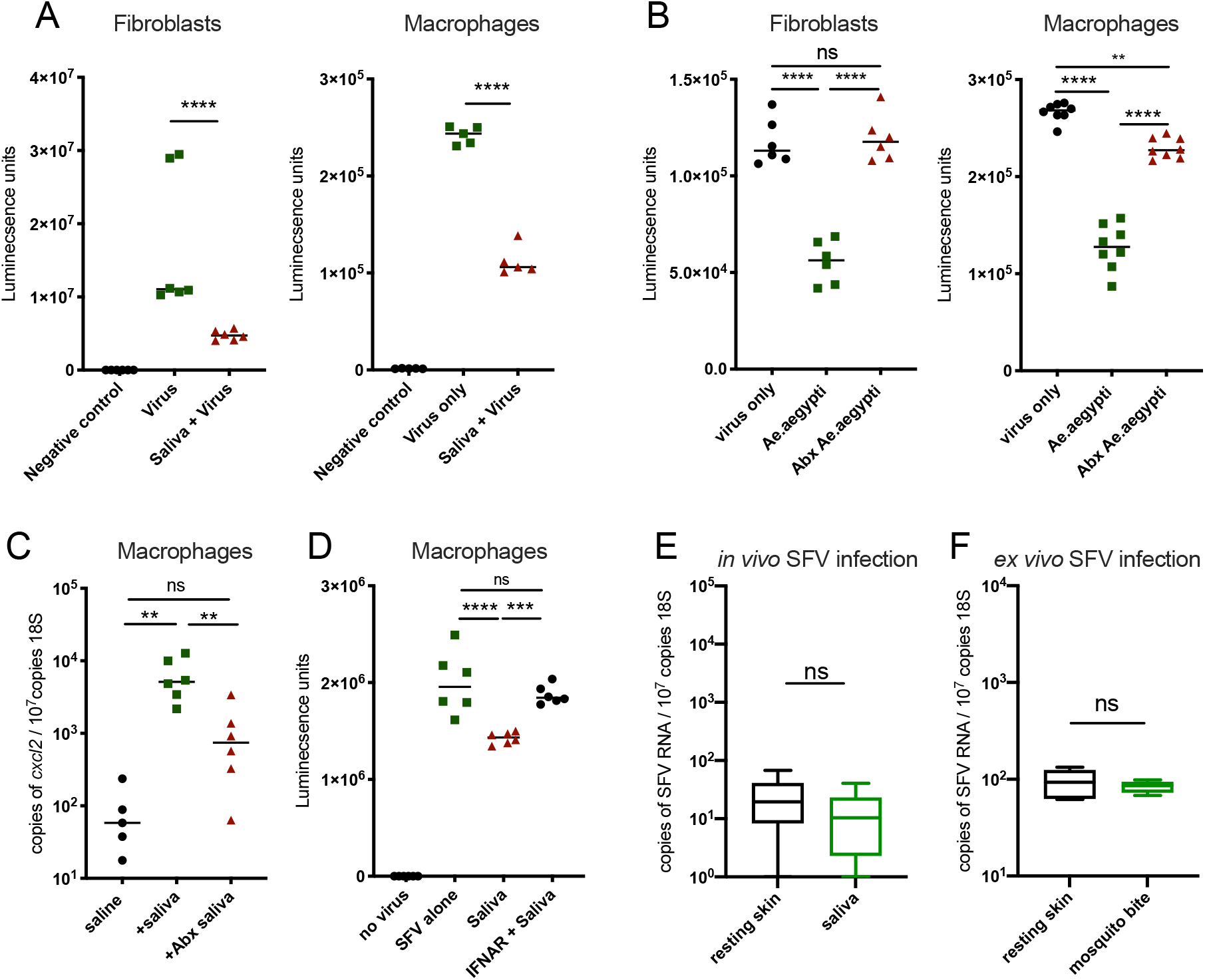
Enhancement of virus infection in vivo requires processes that are absent ex vivo. (A-D) Primary cultures of macrophages and dermal fibroblasts (both principal targets of SFV in vivo) were infected with Gluc-expressing SFV at an MOI of 0.1 and luciferase activity of tissue culture supernatant assayed at 6 hpi. Cells were exposed to virus with or without 0.67μg of saliva protein per well. Cells were treated with saliva for 1 hour prior to infection at room temperature. (A) Cells were exposed to virus with or without saliva (n=5). (B) Cells infected with SFV alone or with saliva from untreated or antibiotic (Abx) treated Ae.aegypti mosquitoes (n=8 and n=6). (C) Macrophages treated with saliva from untreated or Abx-treated Ae.aegypti mosquitoes. Expression of cxcl2 transcripts were measured by qPCR at 6h post treatment (n=6). (D) Macrophages left resting or pre-treated with IFNAR-1 blocking antibody for 1h and then infected with SFV (MOI=1) alone or with saliva from untreated or Abx-treated mosquitoes (n=6). (E) Mice were culled and skin immediately infected with 105 PFU SFV4 alone or with 1.86μg of mosquito saliva. After 15 minutes, to allow for infection of skin-resident cells, skin was dissected and placed in explant culture for 24h. Viral RNA and host 18S were quantified by qPCR (n=8). (F) Mouse skin was bitten by Aedes mosquitoes and inflammation allowed to develop for 16 hours, then skin biopsies of this site infected ex vivo with 105 PFU SFV and viral RNA and host 18S were quantified by qPCR (n=6).

### A heat-sensitive salivary factor from female mosquitoes enhances virus infection independent of bacterial microbiota

Next, we wanted to define which component in saliva was responsible for enhancing virus infection *in vivo*. Because bacterial microbiota in saliva is inflammatory and some inflammatory responses *in vivo* to mosquito biting can enhance infection with virus (Pingen et al., 2016), we hypothesised that microbiota may account for the ability of saliva to enhance virus infection in mice. However, while microbiota-depleted saliva induced significantly lower quantities of mosquito bite-associated inflammatory gene expression in skin (Fig 3A), this did not alter the ability of saliva to enhance virus infection in mice (Fig 3B). Thus, bacterial microbiota was dispensable for the infection promoting ability of saliva *in vivo*. Instead, we did find that the ability of saliva to enhance infection was sensitive to protein-denaturing temperatures (Fig 3C). Together, this suggested that bacteria, although pro-inflammatory, did not influence the ability of saliva to enhance infection and that induction of these inflammatory cytokines by saliva is not a limiting factor. Thus, the pro-viral factor in saliva is likely a proteinaceous factor expressed and secreted by the mosquito salivary gland.

**Figure 3:**
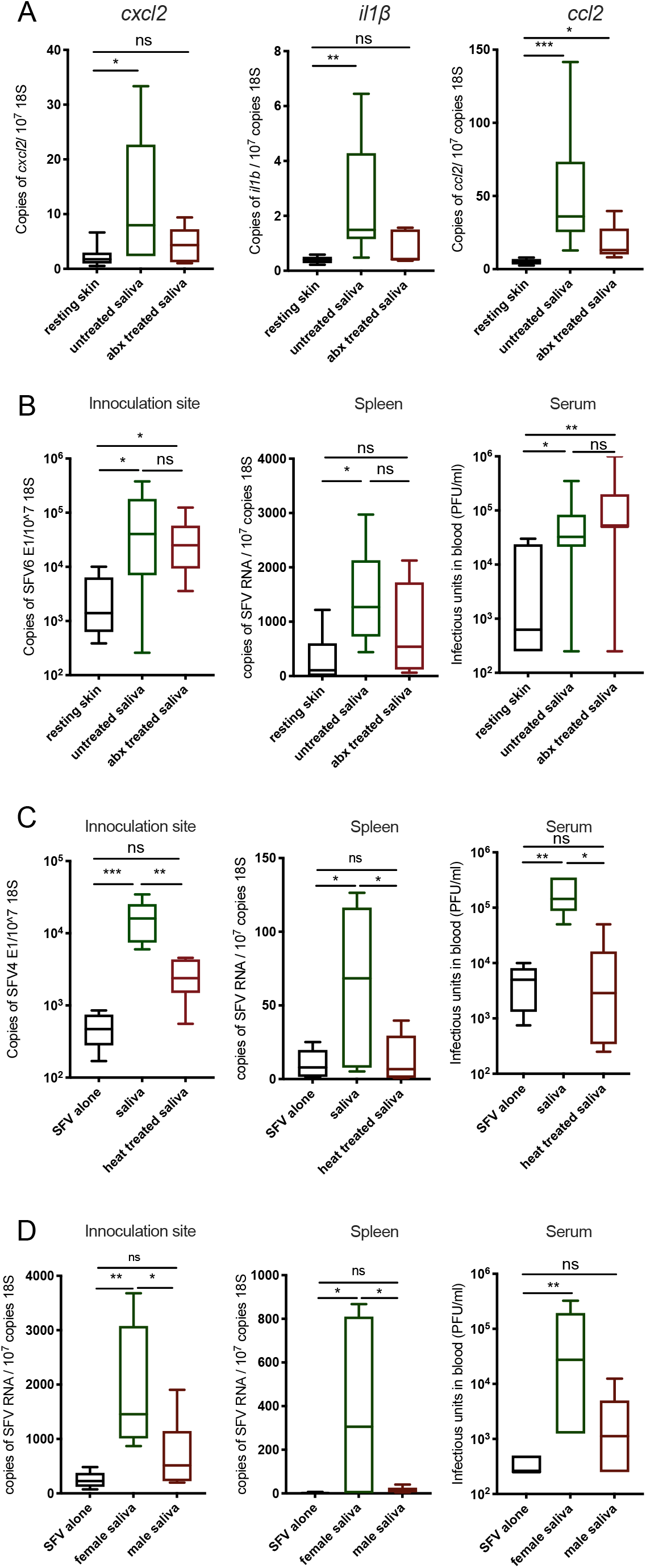
A heat-sensitive salivary factor from female mosquitoes enhances virus infection independent of bacterial microbiota. (A) Mouse skin was injected with 1.86μg saliva from control and Abx-treated mosquitoes and host expression of cxcl2, il1b and ccl2 transcripts assessed at 6 hours (n=6). (B-D) Mouse skin was inoculated with 10^4^ PFU of SFV4 alone or with Ae.aegypti saliva in the upper skin of the left foot. Viral RNA and host 18S were quantified from skin and spleen by qPCR and viral titres of serum by plaque assays at 24hpi. (B) Female Ae.aegypti saliva from Abx-treated or untreated mosquitoes (n>6). (C) Heat treated (10min at 95°C) or untreated female Ae.aegypti saliva (n=6). (D) Male or female Ae.aegypti saliva pooled from 5 mosquitoes combined, reared in the same cage (n=6).

Many female-specific mosquito salivary components have evolved to facilitate blood feeding. In contrast, male mosquitoes do not bite or blood feed and therefore their saliva is naturally deficient in blood feeding factors (Ribeiro et al., 2016). Hence, to determine whether the virus-enhancing salivary factor is one of these factors, we compared the ability of male and female saliva to modulate host susceptibility to virus. Male and female saliva were obtained from mosquitoes caged and reared together. Crucially, we found that female *Aedes* saliva enhanced virus infection to a significantly higher extent than male saliva for saliva. This was the case when either equal volumes of male/female saliva were used (Fig 3D), or when saliva quantities were normalised for protein content, which were substantially lower in male saliva isolates (sFig 3A). Because blood feeding causes changes in gene expression within the salivary glands (Thangamani and Wikel, 2009) we additionally assessed how saliva acquired from previously blood fed and exclusively sugar fed *Ae.aegypti* females modulated virus infection in mice. However, saliva from either group possessed similar virus-enhancing properties, suggesting that a factor that is expressed irrespective of prior blood-feeding was responsible (sFig 3B). Together, this suggests that a female-specific salivary factor, which may have evolved to support efficient blood feeding, was responsible for enhancing susceptibility to virus infection.

### *Anopheles* mosquito saliva lacks the ability to enhance virus infection

To help us identity the virus-enhancing factor present in *Ae.aegypti* mosquito saliva, we next determined whether saliva from other mammalian blood-feeding mosquito species have similar virus infection-enhancing priorities. Saliva from several mosquito species can enhance the infection of viruses that they are competent to transmit. This includes the ability of *Ae.aegypti* saliva and/or bites to enhance infection to a wide range of genetically distinct arboviruses, while *Culex tarsalis* saliva also enhances WNV (Styer et al., 2011). Importantly however, it is not clear whether enhancement by saliva from mosquito species belonging to *Aedes* and *Culex* genera (*Culicinae* subfamily, *Culicidae*), which are also important arbovirus vectors, are comparable in potency. Neither is it clear if this is a general phenomenon elicited by all blood-feeding mosquito species saliva, or whether saliva from arbovirus vector-incompetent mosquitoes such as *Anopheles* (*Anophelinae* subfamily, *Culicidae*) can also enhance infection. Therefore, we compared the ability of saliva from genetically-distinct mosquito species to enhance infection in mice with our two model viruses, SFV and ZIKV, thus enabling a quantitative comparison of mosquito species-specific saliva. Normalised by protein content, *Ae.albopictus* and *Cx.pipiens* saliva enhanced infection to a similar extent as *Ae.aegypti* saliva (sFig 4A,B). However, saliva from *Anopheles* species mosquitoes (*An.gambiae* and *An.stephensi)* uniquely lacked the ability to modulate host susceptibility to infection with either SFV or ZIKV at all time points (Fig 4A-C sFig 4C-E). Furthermore, unlike *Ae.aegypti* bitten skin, *An.gambiae* bitten skin exhibited similar host susceptibility to infection as resting skin (sFig 4C). Thus, while both *Aedes* and *Anopheles* saliva have evolved factors to facilitate efficient blood feeding (Ribeiro et al., 2010), they differ in their ability to enhance infection of mammals with virus.

**Figure 4:**
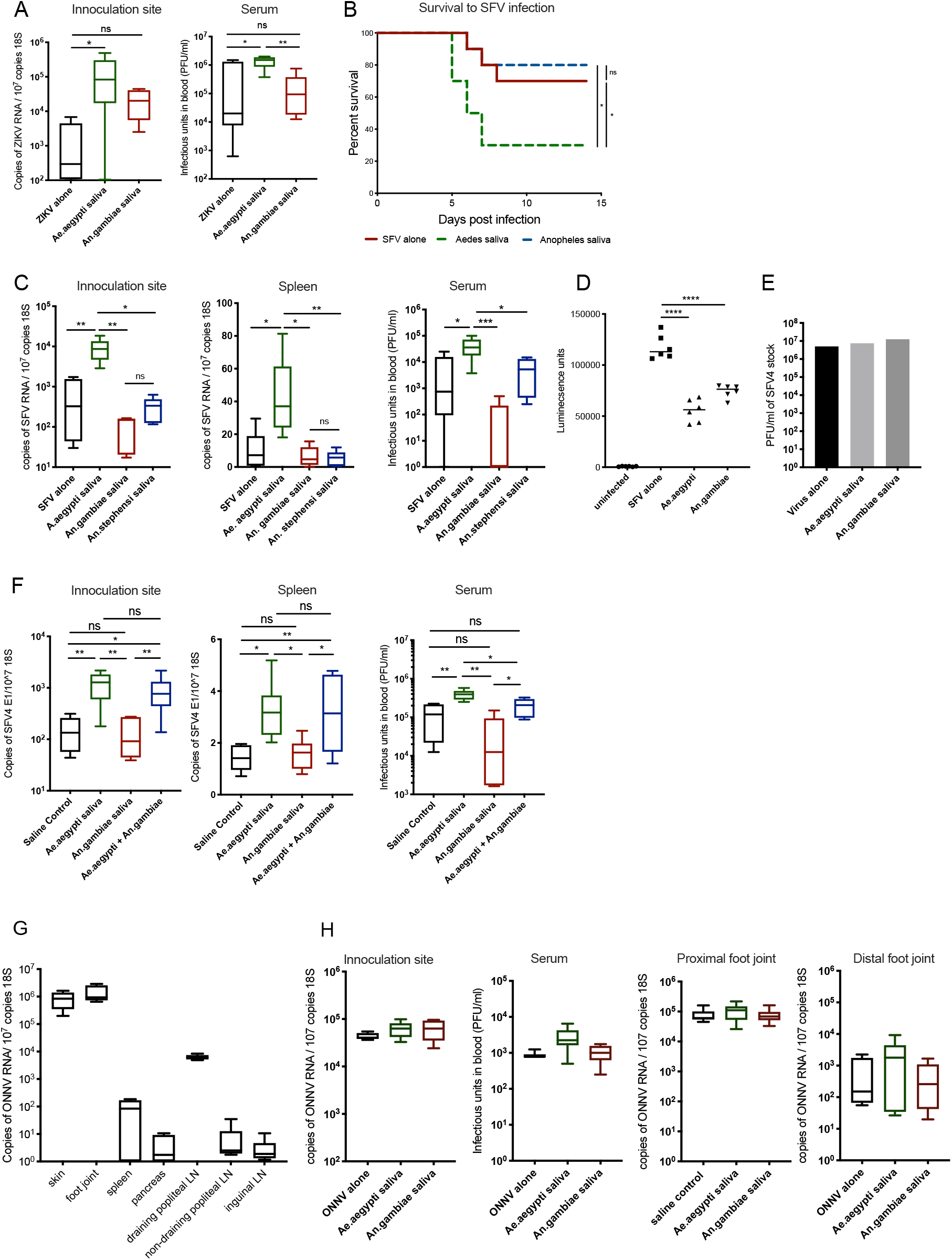
Anopheles mosquito saliva lacks the ability to enhance virus infection. (A-C) Mouse skin was inoculated with either 10^5^ PFU of ZIKV, or 10^4^ PFU of SFV4 alone or with 1.86μg saliva of either Ae.aegypti, An.gambiae or An.stephensi. Virus RNA and host 18S and serum viral titres were quantified at 24 hpi (B) Survival of mice infected with 4×10^4^ PFU of SFV4 (D) Macrophages where infected with luciferase expressing SFV at an MOI of 0.1 alone or with 0.66μg of protein of Ae.aegypti, An.gambiae or Ae.albpictus saliva. Luciferase activity of tissue culture media was assayed at 6 hpi (n=6). (E) BHK cells were infected with 10-fold serial dilutions of SFV4 ranging between 25 000 PFU and 0.25 PFU alone or with Ae.aegypti or An.gambiae saliva and then immediately overlayed with avicel. PFU were then assessed at 48 hpi. Shown here representative PFU for wells in which plaques were quantifiable. (F) Mouse skin was inoculated with 10^4^ PFU of SFV4 alone or alongside 1.86μg saliva of Ae.aegypti, An.gambiae or both species saliva. Virus RNA and host 18S were quantified from skin and spleen by qPCR and viral titres of serum by plaque assays at 24hpi. (G,H) Mice were treated with 1.5mg anti-IFNAR antibodies (clone MAR1-5A3) and 24 hours later infected with 2 ×10^5^ PFU ONNV s.c. in the skin (upper side of the left foot). (G) ONNV RNA quantities in tissues at 48 hpi were defined by qPCR to define tissue tropism. (H) Mouse skin was infected with ONNV alone or alongside 1.86 μg saliva of either Ae.aegypti or An.gambiae. ONNV RNA and host 18S from tissues were quantified by qPCR and serum viral titres were quantified via plaque assays at 48 hpi.

To exclude the possibility that *Anopheles* saliva may contain a unique factor that inhibited virus infectivity, we assessed whether saliva modulated virus infection *in vitro*. Similar to *Aedes* saliva, *Anopheles* saliva possessed some limited ability to protect cells from infection, likely as a consequence of the microbiota as these mosquitoes were reared in the same conditions (Fig. 4D). However, in BHK-21 cells that lack IFN-signalling, both species’ saliva had no effect on ability of virus to replicate and form plaques (Fig. 4E). In addition, we found that by mixing *Aedes* and *Anopheles* saliva together, we were able to restore enhancement of virus infection *in vivo* (Fig 4F). Together, this suggests that the ability to enhance virus infection is mosquito species specific and that the factor responsible in *Aedes* sp. saliva is missing in *Anopheles* saliva.

In comparison to *Aedes* mosquitoes, the widely-distributed group of *Anopheles* species mosquito are not able to efficiently transmit arboviruses, despite them sharing an ecological niche that overlaps with arbovirus endemic areas (Mathiot et al., 1990; Nanfack Minkeu and Vernick, 2018). The sole exception to this is the *Anopheles*-transmitted O’nyong’nyong virus (ONNV), whose genetic sequence is highly similar to CHIKV and similarly disseminates to joints and can cause arthritis (Vanlandingham et al., 2005; Williams et al., 1965). Both *Aedes* and *Anopheles* share some homology in their salivary gland transcriptome (Das et al., 2010; Ribeiro et al., 2007) and both have evolved salivary factors that facilitate efficient blood-feeding (Ribeiro et al., 2010). It is not clear why *Anopheles* sp. mosquitoes are such poor vectors of virus, although there are likely multiple reasons (Vanlandingham et al., 2005). It is conceivable that one reason is the absence of salivary factors that enhance infection of virus in the mammals they feed on. To help explore this, we investigated whether *Anopheles* saliva, or indeed *Aedes* saliva, could modulate infection of mice with ONNV, utilising a newly developed ONNV mouse model in which virus efficiently replicated and disseminated from skin inoculation site to joint tissue (Fig 4G). Importantly, neither mosquito species saliva (Fig. 4H), or indeed presence of mosquito bite (sFig 4F), could modulate host susceptibility to infection with ONNV. Instead, ONNV was able to infect mice robustly, induce viremia and disseminate to joint tissue without the need for saliva-based enhancement of infection. We tentatively suggest that this may partly reflect the absence of virus infection-enhancing factor(s) present in its natural *anopheline* vector.

### Immune sensing of mosquito saliva alone is not sufficient to enhance virus infection

*Aedes* mosquito biting induces the recruitment of monocytic cells, which can be infected by, and replicate within, virus (Pingen et al., 2017, 2016). We made use of our finding above that *Anopheles* and *Aedes* saliva differ in their ability to enhance virus infection to define whether and how *Aedes* saliva activates the recruitment of virus permissive monocytic cells. Firstly, we hypothesised that *Anopheles* and *Aedes* saliva differ in their potency for activating key pro-inflammatory responses that attract monocytic cells and neutrophils. However, we found induction of most inflammatory responses to *Aedes* and *Anopheles* saliva were similar at the transcript level, including the key monocyte chemoattractant *ccl2*, suggesting that innate immune sensing for both was broadly analogous (Fig. 5A). *Anopheles* saliva more potently upregulated some IFN-stimulated genes (ISGs), including *rsad2* and *ccl5* (Fig 5A,B). However, more potent induction of antiviral IFNs by saliva did not account for its ability to modulate virus infection, as enhancement of both SFV and ZIKV infection by *Aedes* saliva was IFN-independent (Fig 5C and Fig 4A respectively). Importantly, we found that despite similar ccl2 chemokine expression, *Aedes* saliva resulted in a robust and rapid influx of CD11b+CD45+Ly6C+ monocytic cells by 2 hours, while *Anopheles* saliva did not (Fig 5D). This did not apply to other myeloid cells, as despite a robust induction of *cxcl2* expression, we saw no influx of neutrophils in response to either mosquito species’ saliva at this early timepoint. Thus, the early expression chemokine induced by saliva was not sufficient on its own to induce a rapid monocytic influx.

**Figure 5:**
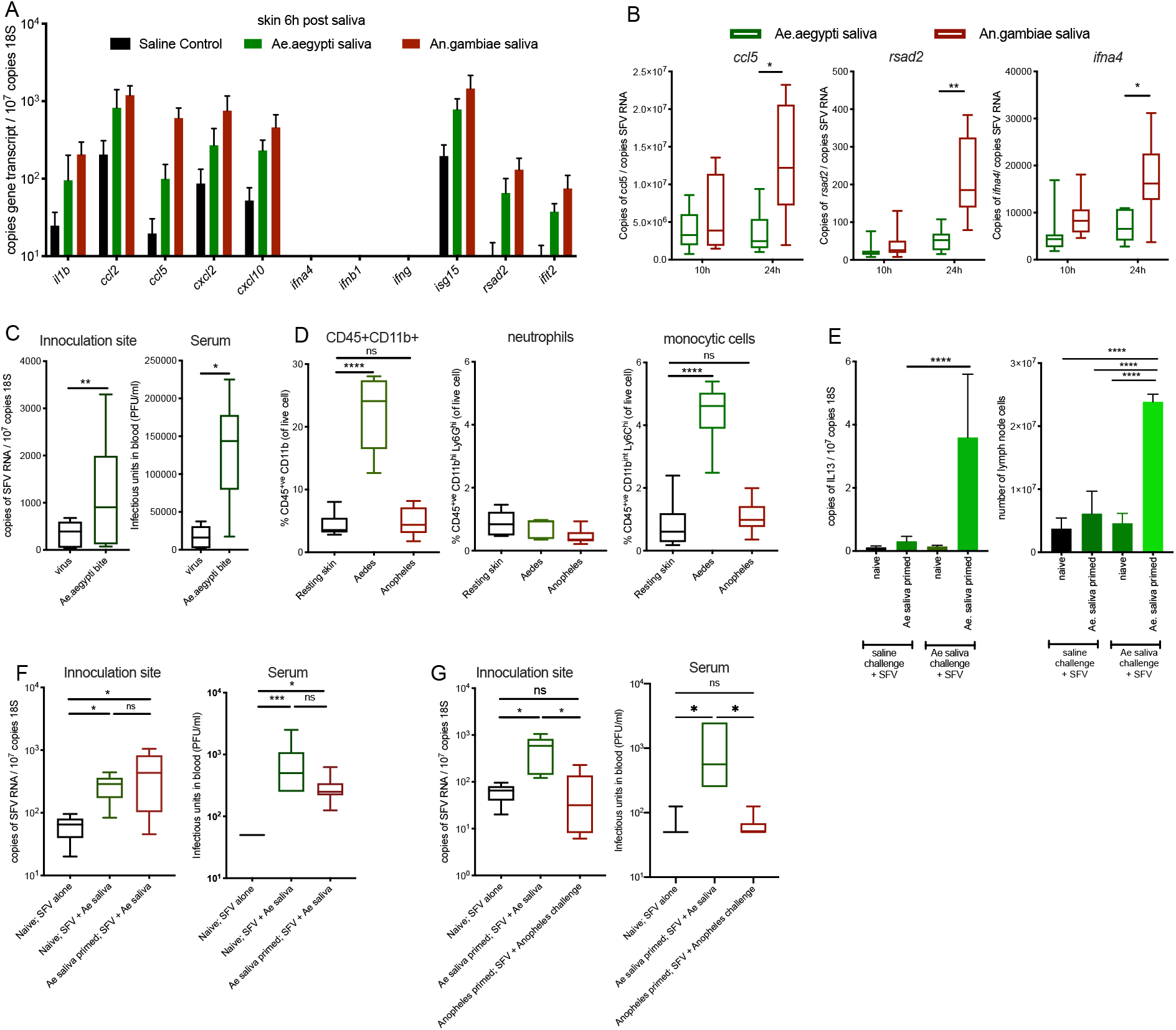
Immune sensing of Aedes saliva is not sufficient to enhance virus infection. (A-B) Mouse skin was injected with either saline control or 1.86μg saliva of either Ae.aegypti or An.gambiae. (A) Copy number of host transcripts in the skin was determined by qPCR at 6 hours (n=6) (B) Mice were inoculated with 10^4^ PFU of SFV4 with either Ae.aegypti or An.gambiae saliva and transcripts quantified by qPCR (n=8). (C) Mice treated with IFNAR-1 blocking antibody a day prior to inoculation with 10,000 PFU SFV4 into mouse skin (resting or following mosquito biting). Viral RNA and host 18S were quantified from skin and spleen by qPCR and viral titres of serum by plaque assays at 24hpi. (n=6) (D) Mosquito-bitten mouse skin was injected with 5 mosquitoes worth of either Ae.aegypti or An.gambiae saliva. At 2 hours, skin from the inoculation site was biopsied and digested to release cells and numbers of myeloid cells (CD45+CD11b+), neutrophils (CD45+CD11b+Ly6G+Ly6C^int^) and myelomonocytic cells (CD45+ CD11b+ Ly6G- Ly6C+ cells) quantified (n=6) (E-G) Balb/c mice skin was inoculated with 10,000 PFU of SFV alone or with saliva. Mice were either naïve to saliva or primed to saliva by prior injections of mosquito saliva weekly for 4 consecutive weeks. (E) IL-13 transcript expression and cell numbers of draining popliteal lymph nodes at 2 hpi. IL-13 transcripts were quantified by qPCR (n=6). (F) Virus RNA in skin was measured by qPCR and serum virus quantified by plaque assay, at 24 hpi (n=6) for mice primed with either Aedes saliva or saline alone. (G) Mice were pre-sensitised to either Aedes or Anopheles saliva for 4 weeks prior to SFV infection co-injected with respective species saliva. Expression of viral SFV gene was measured using qPCR in the skin and serum virus quantified by plaque assay at 24 hpi.

We also found that enhancement of inflammatory responses to saliva, in mice ‘vaccinated’ with saliva prior to infection, did not result in any further modulation of SFV infection. This is relevant as adaptive immune responses that heighten skin inflammatory responses to saliva can occur in those previously exposed to biting mosquitoes. Here, mice were immune sensitised to saliva or saline control by four weekly injections and then on week five injected with either SFV alone or SFV with saliva. At 2 hpi, saliva primed mice exhibited significantly elevated dLN cellularity, IL-5 and IL-13 expression and increased serum IgE indicative of immune-sensitivity to saliva (Fig 5E and sFig 5). However, *Aedes* saliva primed mice showed no difference in their susceptibility to SFV when co-compared to saliva naïve mice (Fig 5F). Importantly, *Anopheles* saliva-primed mice also exhibited no difference in susceptibility to virus, such that they were similar to saliva-naïve mice injected with virus alone (Fig 5G). Together, this demonstrates that immune responses to mosquito saliva per se are not sufficient to enhance host susceptibility to virus.

### Increased vascular permeability induced by *Aedes* saliva enhances virus infection

During the course of our above described experiments, we anecdotally noted that during dissection, mice given *Aedes* saliva accumulated fluid at site of inoculation, which was absent in *Anopheles* saliva injected mice. We hypothesised that *Aedes* saliva may activate the blood vasculature to alter barrier function, enabling the dysfunctional rapid influx of monocytic cells (Fig 5D), and that altered blood vascular permeability might underpin the ability of saliva to enhance virus infection. We therefore quantified levels of oedema as a measure for vasculature barrier leakage following injection with saliva or exposure to mosquito biting. Importantly, both *Ae.aegypti* saliva and bites led to significantly more oedema than *An.gambiae* saliva or biting (Fig 6A, 6B and sFig 6A). Indeed, the complete lack of oedema following *Anopheles* saliva injection suggested it lacked a key factor capable of modulating tissue fluid flow into skin.

**Figure 6:**
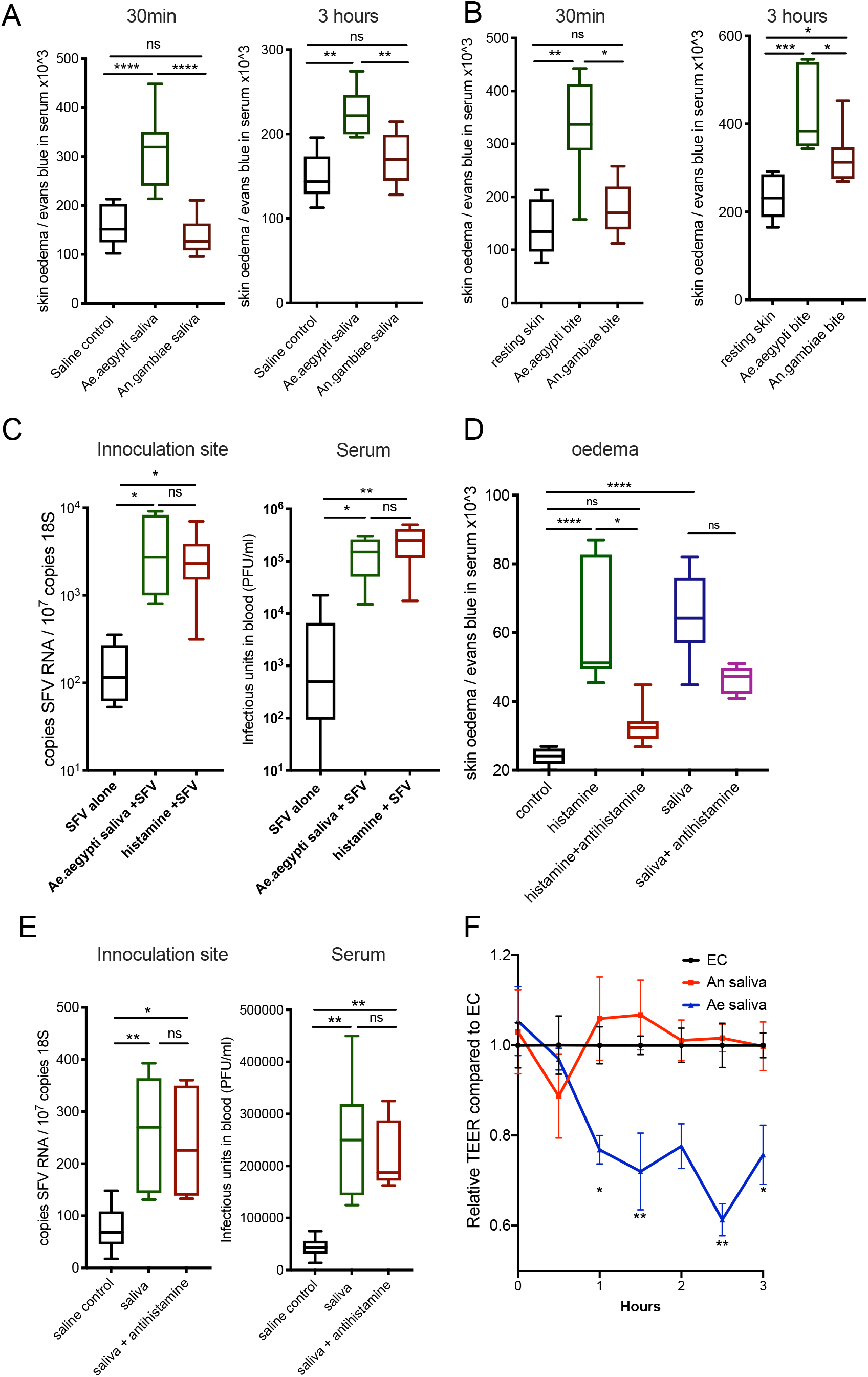
Increased vascular permeability induced by *Aedes* saliva enhances virus infection. (A,B) Mice administered i.p. with Evans blue, were injected with 1.86μg of mosquito saliva in the skin, or exposed to up to 3 bites from Ae.aegypti or An.gambiae and extent of oedema assessed by quantification of Evan’s blue dye leakage into skin at 30min and 3h post saliva/biting via colorimetric assay (n=6). (C) Mouse skin was inoculated with 10^4^ PFU of SFV4 alone or with either Ae.aegypti saliva, or 10 μg histamine dihydrochloride. SFV RNA and host 18S and serum viral titres were quantified at 24 hpi. (D) Mice were administered i.p with Evans blue and then given either control or antihistamines s.c.(0.5mg Cetirizine in 100μl, 0.02mg Loratadine in 100μl and 0.1mg of Fexofenadine in 200μl) 1h prior to saliva injection. Mouse skin was then injected with 5 mosquitoes worth of Ae.aegypti saliva, 10 μg histamine dihydrochloride, or An.gambiae saliva, or a combination of An.gambiae saliva and 10 μg histamine. Quantity of skin Evans blue was measured after 30min by colorimetric assay. (E) Mice were pre-treated with either control saline or antihistamines s.c (0.5mg Cetirizine in 100μl, 0.02mg Loratadine in 100μl and 0.1mg of Fexofenadine in 200μl) 1 hour prior to infection and then skin inoculated with 10^4^ PFU of SFV4 alone or with Ae.aegypti saliva. SFV RNA and host 18S and serum viral titres were quantified at 24 hpi. (F) Human primary endothelial cell monolayers were treated with either control saline, Ae.aegypti or An.gambiae and electrical resistance across the monolayer assessed longitudinally.

We next asked whether induction of vascular leakage was immune-mediated or due to direct action of salivary factors on the vasculature itself. Histamine is the principle host factor that mediates early/immediate oedema in response to exogenous agents, infection or injury (Thangam et al., 2018). Interestingly, we found that injection of exogenous oedema-inducing histamine alone made mice more susceptible to virus infection in the absence of any mosquito salivary factors (Fig 6C), suggesting that endothelial leakage alone is sufficient to promote virus infection. However, we found that histamine was not necessary for host responses to saliva, as treatment with antihistamines did not significantly suppress saliva-induced oedema (although control histamine-induced oedema was suppressed), or virus enhancement by saliva (Fig 6D,E). Together with our findings above (Fig 5E) that immune hypersensitisation to saliva did not modulate host susceptibility to virus, we hypothesized that the *Aedes* salivary factor acts directly on blood vasculature, in an immune response-independent manner. Therefore we exposed cultured endothelial cell monolayers to saliva and defined their barrier function over time. Crucially, we found that *Aedes* saliva, but not *Anopheles* saliva, disrupted endothelial barrier function within an hour, similar to that observed *in vivo* (Fig 6F).

Collectively, by comparing host responses to pro-viral *Aedes* saliva and *Anopheles* saliva we have identified blood vascular leakage as a key feature associated with female *Aedes* saliva enhancement of virus, a phenotype that is recapitulated with histamine-induced barrier loss, but which is independent of innate and adaptive immune sensing of saliva. Instead we found a salivary factor in *Aedes* saliva was responsible for directly inducing endothelial barrier loss.

### *Aedes* sialokinin is sufficient to induce blood vasculature barrier leakage and enhance virus infection

The saliva of blood-feeding arthropods contains many factors that increases the volume of blood reaching the arthropod mouthparts, including those that may alter vascular function. In *Ae.aegypti*, one such hypothesised factor is sialokinin (SK), a 1400-Da peptide product of the pro-sialokinin precursor-encoding gene (AAEL000229). SK belongs to the family of tachykinin-like peptides (Ribeiro, 1992; Champagne and Ribeiro, 1994) and is expressed in the female salivary gland (Beerntsen et al., 1999). We therefore sought to define the evolutionary conservation of homologues in other blood-feeding insects and determine whether SK has any role in modulating mammalian vascular barrier function and thereby host susceptibility to virus infection in mice.

Tachykinin-related peptides (TRPs) have been identified from many insect species (Yeoh et al., 2017) and in *Ae.aegypti* five TRPs are encoded by Tachykinin (AAEL006644), a gene with single-copy orthologues across protostome invertebrates (Broeck et al., 1999; Nässel et al., 2019). Intriguingly, we found that the SK tachykinin-like peptide (NTGDKFYGLMamide) more closely resembles typical deuterostome-type (FXGLMamide) than protostome-type (FXGXRamide) peptides (Fig 7A). In addition, SK is also recognised by the PROSITE tachykinin family signature pattern (PS00267), which matches deuterostome-type but not protostome-type peptides. Together, this suggests that, being more similar to deuterostome-type tachykinins, it might primarily target exogenous vertebrate receptors rather than endogenous mosquito receptors. Indeed, SK is able to elicit intestinal contraction at levels similar to the prototypic tachykinin, mammalian substance P, as well as cross-desensitization with substance P and reaction with an anti-substance P antibody (Ribeiro 1992, Champagne and Ribeiro 1994). The presence of such deuterostome-type tachykinin-like peptides in arthropods appears to be extremely rare, with the only matches to the PROSITE pattern being to a peptide from a fungus-growing ant, and four peptides from the highly venomous Brazilian wandering spider. Importantly, this venom, like *Ae.aegypti* saliva, induces extensive oedema in mammals (Palframan et al., 1996). Beyond arthropods, four other peptides matching the sequence were identified in three octopus species, where vasoactive effects might be also important for immobilising vertebrate prey. Thus, the SK peptide sequence resembles both vertebrate tachykinin and a spider venom peptide that can induce potent inflammatory responses in mammals, including oedema.

**Figure 7:**
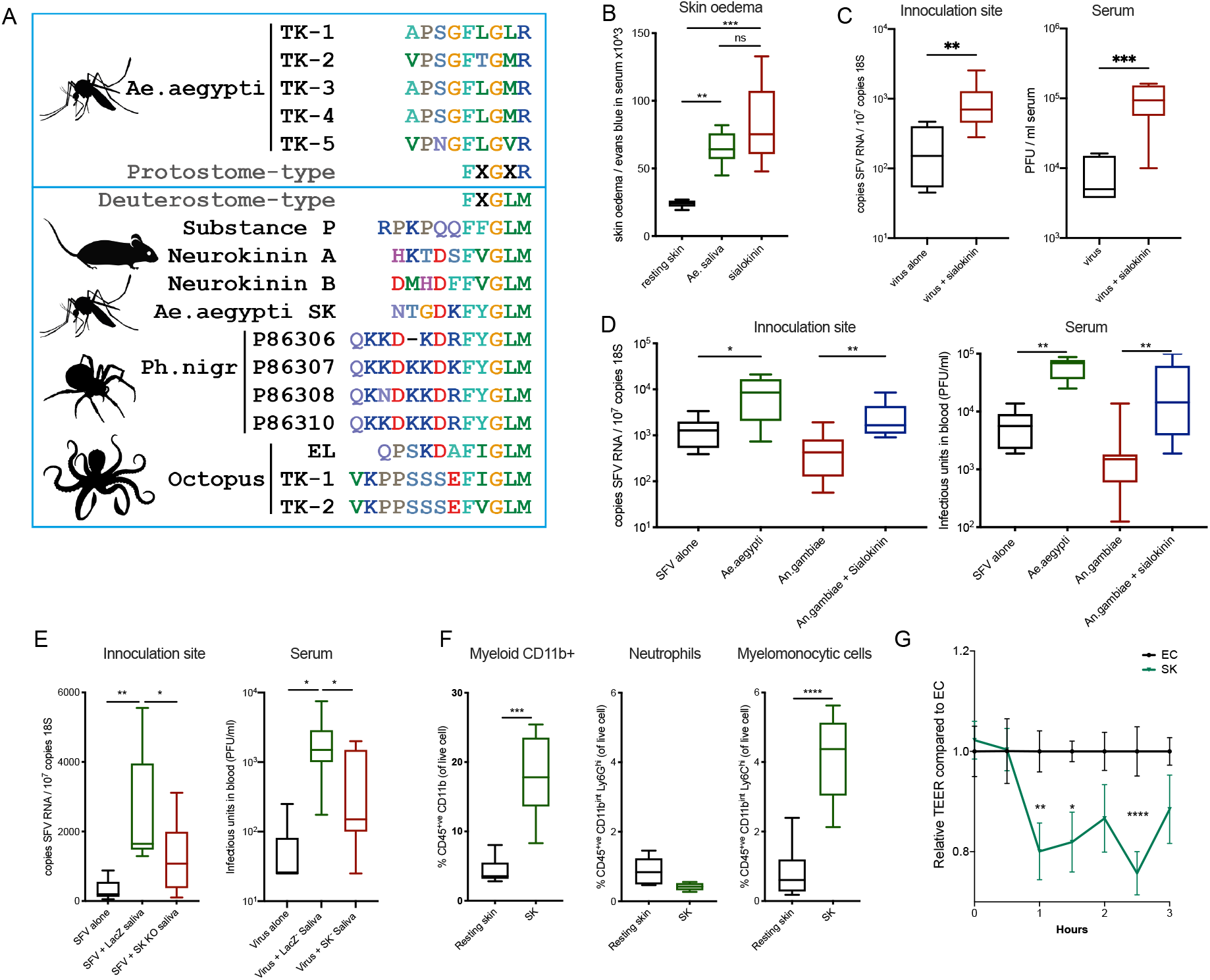
*Aedes* sialokinin is sufficient to induce blood vasculature barrier leakage and enhance virus infection. (A) Protostome and deuterostome-type tachykinin-related peptides. The Aedes aegypti Tachykinin (AAEL006644) gene encodes five tachykinin-related peptides (Ae.aegypti TK1-5) that match the protostome-type FXGXRamide signature (upper box). In contrast, the sialokinin tachykinin-like peptide (Ae.aegypti SK, 1400 Da) matches the deuterostome-type FXGLMamide signature (lower box). This motif is similar to mammalian tachykinins Substance P and Neurokinins A and B. Rare examples of other protostome species with peptides matching the deuterostome-type signature include octopuses (EL, eledoisin; TK, tachykinin) and a venomous spider (Ph.nigr). (B) Mice were injected IP with Evans blue and 1 h later skin injected with either 1.86 μg Ae.aegypti saliva or 1 μg sialokinin. Skin samples were collected 30min pi. (C to E) Mouse skin was inoculated with 10^4^ PFU of SFV4 alone or with saliva from either Ae.aegypti or An.gambiae, with or without supplementation with 1μg sialokinin peptide. SFV RNA and host 18S were quantified by qPCR and serum viral titres quantified by plaque assay at 24 hpi. (E) Virus administered alone or alongside saliva from dsRNA LacZ or dsRNA SK injected Ae.aegypti females. (F) Resting mouse skin was injected with sialokinin alone. At 2 hours, skin from the inoculation site was biopsied and digested to release cells and numbers of myeloid cells (CD45+CD11b+), neutrophils (CD45+CD11b+Ly6G+Ly6C^int^) and monocytic cells (CD45+ CD11b+ Ly6G- Ly6C+) quantified (n=6) (G) Human primary endothelial cell monolayers were treated with either control saline or 1μM SK alone and electrical resistance across the monolayer assessed longitudinally.

To complement the peptide searches, we thoroughly examined the available insect genomics data to define specifies specificity of SK. Here, we performed sequence searches with the pro-sialokinin precursor protein (85 amino acids) and gene (398 basepairs). No homologues were identified from the gene family data at VectorBase (Giraldo-Calderón et al., 2015) or at OrthoDB (Kriventseva et al., 2019). Sequence searches of the National Centre for Biotechnology Information non-redundant protein and nucleotide databases returned hits only to the *Ae.aegypti* pro-sialokinin precursor itself. We also searched the newly available genome assembly for *Culex tarsalis* (Main et al., 2020), which returned no significant hits. Further sequence searches of all nucleotide data at VectorBase (25 mosquito species, genomes, transcriptomes, cDNAs, ESTs) only recovered the *Ae.aegypti* pro-sialokinin precursor itself (also in the *Ae.aegypti* Aag2 cell line genome assembly). The large sampling of *Anopheles* species, including several high-quality chromosome-level assemblies, lends confidence to the conclusion that this gene is not present in anophelines. The lack of hits in *Ae.albopictus* and two *Culex* species suggest that it could be specific to *Ae.aegypti.* However, the gene is located in a repeat-rich gene desert (nearest neighbours are 0.48 Mbp and 0.99 Mbp) at the start of *Ae.aegypti* chromosome one. Such repeat-rich regions can be challenging to sequence and assemble, therefore it remains possible that other culicine mosquitoes do possess pro-sialokinin precursor-encoding genes.

Together this suggests that SK, a female salivary-specific gene that is absent in *Anopheles* saliva and has known mammalian tachykinin-like activity may be responsible for the ability of *Aedes* saliva to promote vascular leak and thereby enhance virus infection. Indeed, when co-inoculated with virus in the absence of saliva, synthesised SK caused a rapid induction of oedema (30 min post injection), and crucially also enhanced virus infection to a similar extent as *Aedes* saliva in mice (Fig 7B and C). In addition, when SK was added to *Anopheles* saliva, viral enhancement was successfully generated (Fig 7D). We also found that when the SK gene expression was silenced by siRNA knockdown in adult female *Ae.aegypti* mosquitoes (sFig 7), by injection of a dsRNA targeting the SK gene, their saliva lost some ability to enhance infection compared to dsLacZ-dsRNA injected control females (Fig 7E). The extent of knockdown of SK gene expression (7-fold reduction of median, sFig 7) was similar to the magnitude of reduction in virus serum titres by 24 hpi (10-fold reduction, Fig 7E). The reduction in skin virus RNA quantities in SK-deficient saliva injected mice was more modest (1.5-fold reduction, Fig 7E), however this may partially reflect the nature of qPCR assay that detects both residual virus RNA genomes and newly transcribed virus RNA. Importantly, similar to whole *Aedes* saliva, the ability of SK alone to enhance virus infection correlated with its ability to selectively and rapidly recruit monocytic cells *in vivo*, but not neutrophils (Fig 7F). This effect was via direct action on endothelial cells as exposure of monolayers to SK resulted in rapid loss in electrical resistance and thereby an increase in permeability (Fig 7G), similar in kinetics and magnitude as that observed with *Ae.aegypti* saliva (Fig 6F). Thus, we have identified a factor in *Aedes* saliva, sialokinin, which induces blood vascular barrier permeability, leading to oedema and a rapid influx of virus-permissive monocytes that enhances infection with virus.

## Discussion

In this report we have defined how mosquito saliva enhances infection of the vertebrate host with mosquito-borne virus. Inoculation of virus at mosquito bites is an important stage, common to all mosquito-borne virus infections. Mosquito-derived factors in saliva, which are co-injected with virus into the skin during biting, enhance infection by enabling recruiting virus permissive monocytic cells. Replication in these cells provides an early replicative boost for virus that fuels higher viremia and thereby more severe clinical disease. In this report, we have identified one such salivary factor, sialokinin, that is responsible and defined its mechanism through direct action on blood endothelial cells. In doing so, we have defined an important determinant of the clinical outcome to infection at the arthropod/mammalian interface. We suggest this can now inform the development of novel vaccine candidates that target SK and other mosquito salivary factors to reduce host susceptibility to arboviral disease. Such vaccines may have wide applicability, as mosquito-saliva enhancement is widely observed for these viruses.

Our original observation that early enhancement of infection by saliva was only evident *in vivo*, is perhaps now no longer surprising, considering the requirement for a circulatory system that was absent in our *in vitro* and *ex vivo* models. Nonetheless, lack of enhancement *in vitro* was at first somewhat surprising, as a number of previous studies have demonstrated an enhancing effect of mosquito saliva in some cells(Conway et al., 2014b). However, many of these studies utilised salivary gland extract, rather than mosquito saliva itself, which may explain the difference in findings. These data also demonstrate the limitation of exclusively using *in vitro* models that don’t fully replicate all features of virus infection during arthropod biting. Next, we sequentially excluded a role for other salivary components, such as microbiota, and instead identified a role for a female salivary gland specific, heat-denaturing sensitive factor. The ability of female *Aedes* saliva, but not *Anopheles* saliva, to act directly on blood endothelial cells to promote barrier loss, rather than solely through immune sensing, enabled a rational approach in selecting SK as a candidate to study. As such, we showed that SK was sufficient and necessary to induce endothelial barrier permeability and enhance virus infection.

The observation that *Anopheles* saliva does not enhance virus is also interesting considering its implication for understanding of vectorial capacity for arbovirus transmission. Our data suggests that coevolution of arboviruses in competent vectors such as *Aedes,* has led to the utilisation by virus of vector-specific factors to increase their infectiousness, which we suggest will modulate the overall effectiveness of arbovirus transmission by the competent vector, leading to an increase in vectorial capacity and thereby disease burden.

## Acknowledgements

We thank Kamila Fras and Kirby Brown, undergraduate students at the University of Glasgow, who helped with salivation of mosquitoes. Mosquito colonies were kindly supplied by; Prof E. Devaney, University of Glasgow, UK (*Ae. aegypti*); Dr A.-B. Failloux, Institut Pasteur, France (*Ae. albopictus* La Providence INFRAVEC2 line); Dr M. Weill, Université de Montpellier, France (*Cx pipiens quinquefasciatus* SLAB strain); and Dr C. Bourgouin, Institut Pasteur, France (*An. coluzzii* Ngousso strain, referred to as *An. gambiae* in this report, and *An. stephensi* Sda 500 strain). We also thank; the University of Leeds Faculty of Medicine & Health Flow Cytometry and Imaging Facilities; the University of Leeds for funding Daniella Lefteri; the University of Leeds St James’ Biomedical Services and Gillian Cardwell for assistance with mouse studies.

## Author Contributions

Conceptualisation by C.S.M, D.A.L and E.P. D.A.L, E.P. and C.S.M conceived and performed experiments and wrote the manuscript. S.R.B, M.P., S.T., E.F.B, G.G., M.V.L, V.M., and R.W. performed experiments. M.V and A.M. provided reagents. R.W., P.C. S.G. and K.S provided expertise, feedback and edited the manuscript. E.P., R.F. and C.S.M secured funding.

## Star Methods

### Resource Availability

Further information and requests for resources and reagents should be directed to and will be fulfilled by the Lead Contact.

### Materials Availability

This study did not generate new unique reagents.

### Data and Code Availability

This study did not generate/analyze [datasets/code].

### Key resources table

#### Cell lines

Cells were kept at −196°C for long term storage.

BHK-21 cells were used to grow-up virus stock and determining viral titres via plaque assays. BHK-21 cells were cultured at 37°C with 5% CO_2_ in GMEM media supplemented with 10% TPB, 5% FCS and 100 units/ml penicillin and 0.1 mg/ml streptomycin.

*Aedes albopictus* mosquito derived C6/36 cells were used for growing virus stocks. C6/36 cells were cultured at 28°C with no added CO_2_ in L-15 media supplemented with 10% TPB, 10% FCS and 100 units/ml penicillin and 0.1 mg/ml streptomycin.

Mouse Embryonic Fibroblasts from C57BL/6 mice were kept at −196°C for long term storage. MEF cells were cultured at 37°C at 5% CO_2_ in DMEM media supplemented with 10% FCS, 100 units/ml penicillin and 0.1 mg/ml streptomycin and 1% Glutamax in flasks pre-coated with 0.2% gelatine.

Macrophages were extracted from C57BL/6 mouse bone marrow by flushing cells from the femur using a 26-gauge needle with cold PBS. Cells were passed through a 40μm cell and cultured at 37°C at 5% CO_2_ in DMEM/F12 media supplemented with 10ng/ml M-CSF, 10% FCS, 100 units/ml penicillin and 0.1 mg/ml streptomycin , 1% Glutamax and 5mg Gentamycin. 4×10^5^ cells were seeded in 10ml media per sterile plastic petri dish used. 7 days post extraction cells were pooled by and adding 3ml Cellstripper (non-enzymatic cell dissociation solution) to each dish and cells were then gently scraped of the plastic. Cells seeded at a concentration of 2×10^5^ per well in a 24-well plate or at 5×10^4^ per well in a 96-well plate in complete DMEM/F12 media.

#### Virus strains

Semliki Forest virus 4 (SFV4) and Semliki Forest virus 6 (SFV6) stocks were generated from plasmids containing corresponding infectious cDNA (icDNA) sequences (Ulper et al., 2008). SFV6-2SG-GLuc (Gaussia luciferase) is modified SFV6 where sequence encoding Gluc marker is inserted under a duplicated SG promoter positioned at 3’ direction of structural reading frame. ONNV-2SG-ZsGreen represents a modified ONNV of Chad isolate where sequence encoding for ZsGreen is inserted between native and duplicated SG promoters of ONNV. Plasmids containing the icDNAs of SFV4, SFV6, or SFV-6-2SG-GLuc were electroporated into BHK-21 cells to generate infectious virus with 2 pulses at 250V for 0.8S. Plasmid to rescue ONNV-2SG-ZsGreen was linearized first with PmeI (NEB) to prepare it for run-off transcription; RNA was transcribed using MEGAscript SP6 Transcription Kit (Thermo Fisher Scientific) in presence of Ribo m7G Cap Analog (Promega). The RNA was transfected to BHK cells using Lipofectamine (2000) to rescue the virus. Rescued SFV6-2SG-GLuc was aliquoted with cellular debris to allow for improved virus uptake by macrophages *in vitro*. Wild type ZIKV from Recife, Brazil was kindly provided by Prof. Alain Kohl at the university of Glasgow. ZIKV was grown in Vero cells and BHK-21 cells, supernatant was collected then centrifuged to remove cell debris and virus titers were determined by plaque assays on BHK-21 cells. All viruses used *in vivo* were passaged once through C6/36 cells. Supernatant from infected C6/36 cells was collected and infectious virus present in the supernatant was titrated via plaque assay in BHK-21 cells. Viruses were diluted in PBSA to 1×10^7^ PFU/ml

#### Mouse strains

Wild type C57BL/6j mice bred in-house at the SBS at the University of Leeds were used in all *in vivo* experiments unless stated otherwise. Mice were maintained at SBS under specific pathogen free conditions and used between 4 and 12 weeks of age unless stated otherwise. BALB/C mice were purchased from Charles River Laboratories. Mice were age and sex matched in all *in vivo* experiments. All procedures were carried out in accordance with the United Kingdom Home Office regulations under the authority of the appropriate project and personal license.

### Mosquito strains

*Ae.aegypti* (Liverpool strain)
*Ae.albopictus* (La Providence strain)
*Cx.pipiens* (slab strain)
*An.gambiae* (Kisumu strain)
*An.stephensi* (SDA-500 strain)

### Primers used in this study

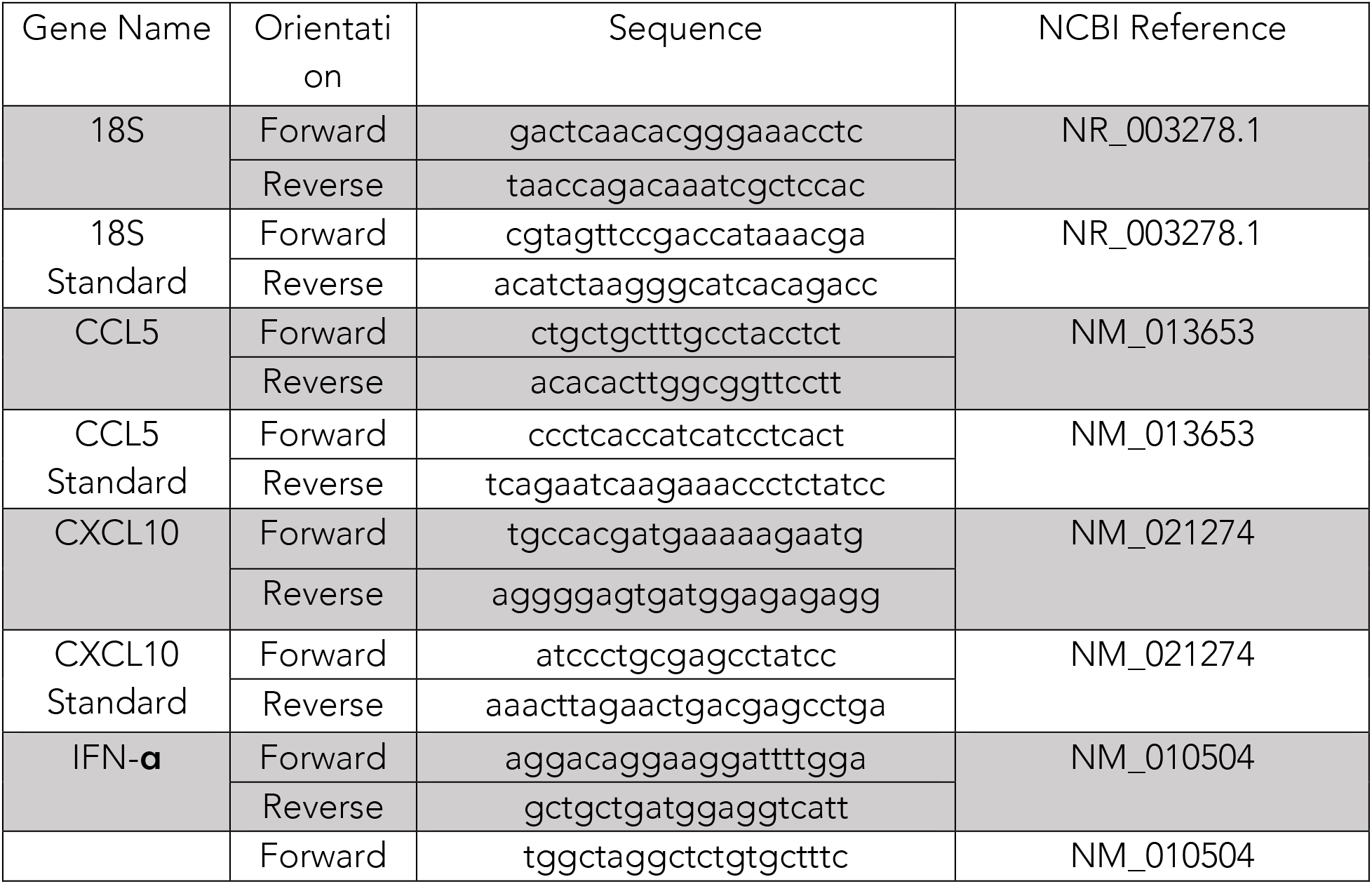
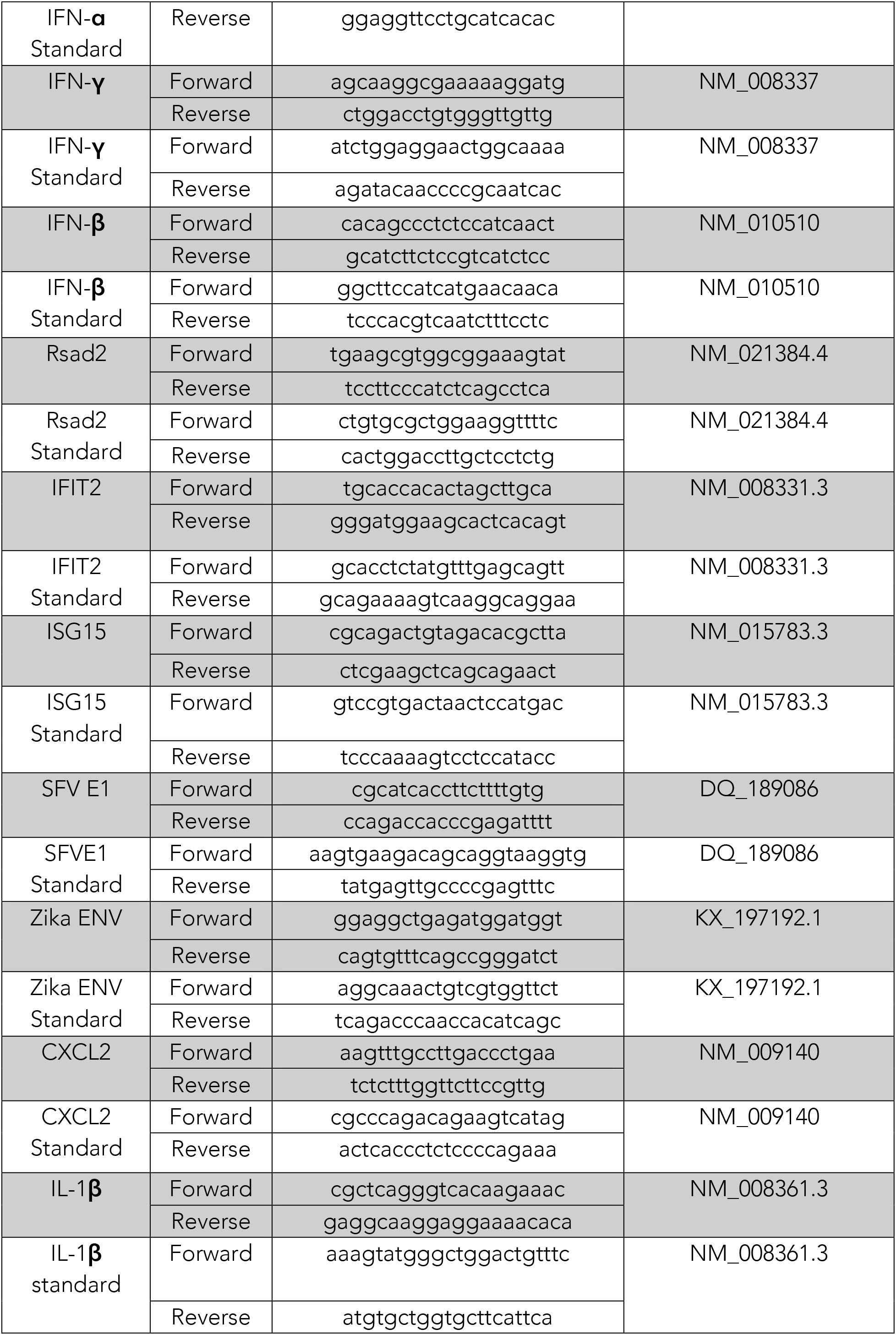
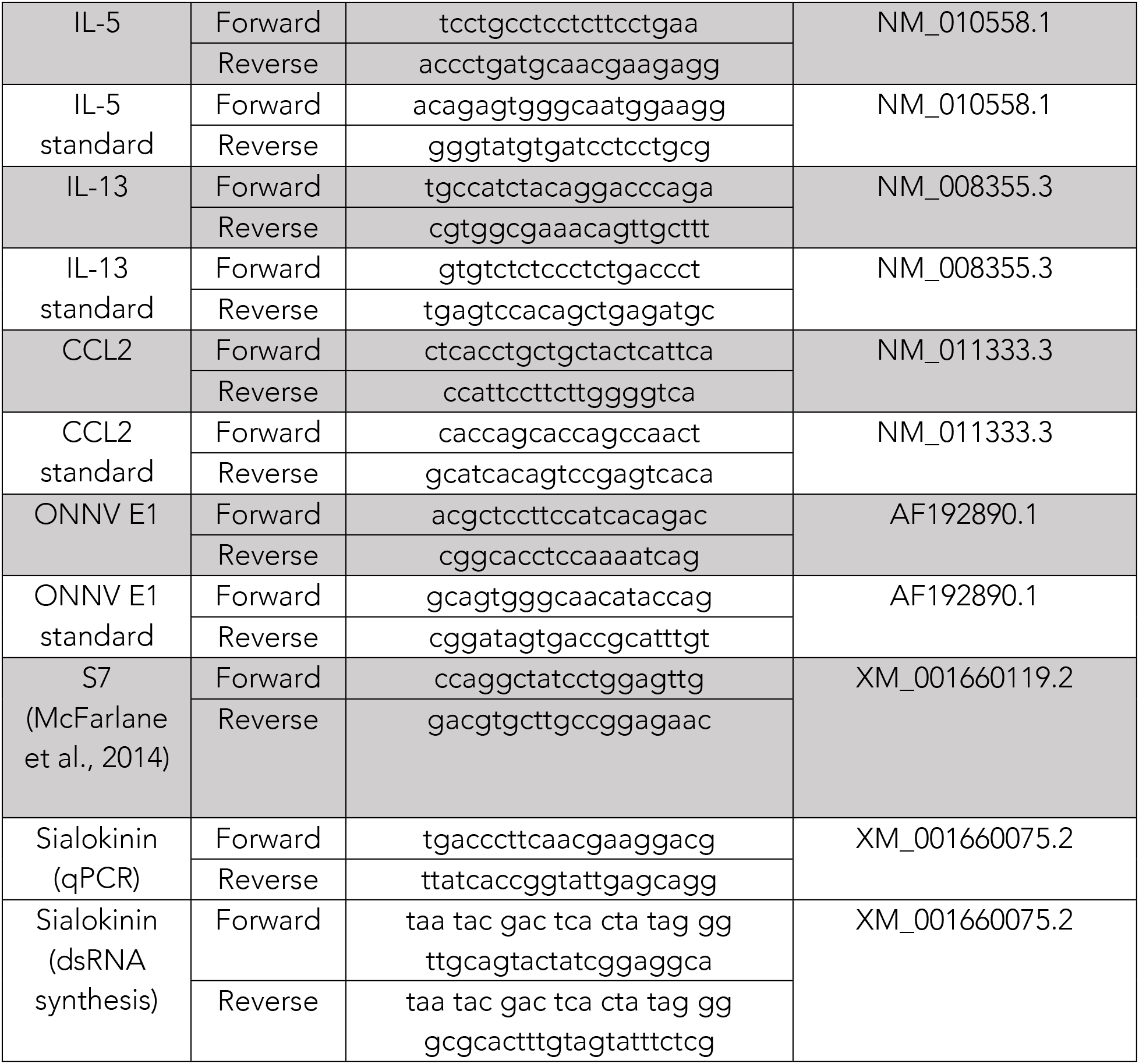

#### Mosquito Rearing and handling

The different mosquito species used were *Ae. aegypti* Liverpool strain (a gift of Prof E. Devaney, University of Glasgow, UK), Ae. albopictus La Providence (INFRAVEC2 line, a gift from Dr A.-B. Failloux, Institut Pasteur, France), Cx pipiens quinquefasciatus SLAB strain (a gift from Dr M. Weill, Université de Montpellier, France), An. coluzzii Ngousso strain and An. stephensi Sda 500 strain (a gift from Dr C. Bourgouin, Institut Pasteur, France). Mosquitoes were reared at 28°C and 80% humidity conditions with a 12 h light/dark cycle. *Ae.aegypti*, *Ae.albopictus* and Cx.*pipiens* eggs on filter paper were placed in trays containing approximately 1.5 cm of water to hatch overnight. Larvae were fed with Go-cat cat food until pupation. *An.gambiae* and *An.stephensi* eggs were hatched the same day of arrival by placing in water. Ground Tetramin fish flakes were fed to the larvae the first days following which the larvae were fed with tetramin pellets until pupation. When pupae formed these were picked and placed in small water filled containers and left to emerge into BugDorm mosquito cages. All adult mosquitoes were fed a 10% sucrose solution. Mosquitoes were ready for salivations and biting experiments 21 days post hatching. Emerging adult mosquitoes were maintained on a 10% (w/vol) sucrose solution ad libitum. Females were fed with heparinised rabbit blood (Envigo, UK) for 1 h using a 37°C Hemotek system (Hemotek Ltd, Blackburn, UK).

For salivation of mosquitoes, the mosquito proboscis was placed in a p10 tip containing 0.5μl immersion oil (Cargille Laboratories, USA). Mosquitoes were then left to salivate for up to an hour before tips were placed in an Eppendorf tube and centrifuged. Saliva droplets were then pooled and stored at −80 °C. Before use, droplets of saliva were carefully pipetted out of the oil under microscope and diluted in PBSA. 5 mosquitoes worth of saliva was utilised per injection unless stated otherwise.

#### Antibiotic treatment

Mosquitoes were given a 10% sugar solution containing a mixture of 200 units/ml penicillin and 0.2 mg/ml streptomycin , Gentamycin at 200 μg/ml Gentamycin and Tetracycline at 100 μg/ml from emergence and for 7 days. Validation of antibiotic treatment was conducted by counting CFU (sFig 1C).

#### Luciferase assay

Luciferase assays were performed with SFV6-2SG-GLuc. Bone marrow derived M-CSF macrophages or MEF cells were seeded at a known concentration in 24 or 96 well plates and infected with a known amount of SFV6-2SG-GLuc. Cells were either pre-treated with mosquito saliva, or saliva was added premixed with the virus. For detection of luciferase in macrophages and fibroblasts infected *in vitro* Renilla Luciferase Assay System (Promega) kit was used and samples were run on Mithras LB 940 Multimode Microplate Reader.

#### Oedema

Oedema was measured by the use of systemically injected Evans Blue dye which binds covalently to serum albumin. During normal physiological conditions, the endothelial cell barriers lining the blood vessels prevent the passage of albumin into tissues. When endothelial barrier function is disrupted however during inflammation, macromolecules such as albumin are able to pass through. Therefore, the measuring of the concentration of Evans Blue dye at the site of inflammation was quantified as indictor of the oedema.

In order to determine the amount of fluid accumulation and vascular leakage in the skin, mice were injected sc with 200μl of 1% Evans Blue. Skin samples were acquired 30min, 3h or 6h post challenge and placed in 250μl of formamide and left to soak overnight at 4°C. Skin samples were then removed from the solution and the dye-stained formamide solution was taken and a 10-fold serial dilution was created by mixing the samples with water. Levels of fluid accumulation was determined using colorimetric measurement of dye concentration at 620nm using the Multiskan EX. Blood samples were acquired and centrifuged. Amount of dye present in the serum was used as a control for amount of dye present in each mouse. In order to ensure the complete removal of any residual dye from the blood in the skin tissue perfusions were carried out immediately after acquiring blood samples. During this process, using a 50ml syringe of PBS with a 26-gauge needle which was inserted into the ventricles and the PBS pumped in to ensure the flush out of blood from the entire circulation.

#### Antihistamine treatment

Antihistamine treatment was performed as follows: 0.5mg Cetirizine in 100μl, 0.02mg Loratadine in 100μl (approx. 1mg/kg) and 0.1mg of Fexofenadine in 200μl (approx. 5mg/kg). Cetirizine and Fexofenadine were pre-mixed and given as a 300μl IP injection whilst Loratadine was given as a separate IP injection of 100μl.

#### *In vivo* mouse infections

Mice were anesthetized with 0.1ml/10g of Sedator/Ketavet IP injection and placed on foil on top of the mosquito cages with the dorsal side of one or both hind feet exposed allowing max 5 mosquitoes to feed. Toes were covered with tape to prevent mosquitoes biting. Mosquitoes left to feed until fully engorged. Virus injections of either C6/36 derived SFV6 (250 PFU in 1μl) or SFV4 (10000 PFU in 1μl) were made directly at the bite site with a 5μl 75N syringe, 33gauge (Hamilton) using small RN ga33/25mm needles (Hamilton). Saliva injections were made at a concentration of 5 mosquitoes-worth of saliva per injection.

Mice were culled via a schedule 1 method. Tissues dissected depended on the experiment but most commonly included, skin from foot and spleen. Blood samples were also collected from the ventricles. Tissue samples collected were stored in 0.5ml RNAlater in 1.5ml tubes, with the exception of spleen and brain samples that were cut in half and stored in 1ml of RNAlater to enable complete penetration of the RNAlater in to the tissue. All samples were left in RNAlater for a minimum of 16 h to prevent RNA degradation. Samples were then stored at 4°C short term storage or at −80°C for long term storage. Blood samples were centrifuged and serum was collected and stored at −80°C until use. Tissue samples were then analysed via qRT-PCR for analysis of the expression of SFV virion glycoprotein E1, which is a gene expressed via SG RNA and that we have previously been established as a good indicator of total viral RNA levels (genome plus transcripts). Serum was analysed for viral titres via plaque assays.

#### Survival

Mice subjected to neurotropic virus infections were monitored 4 times daily and weighed every morning for the entire duration of the experiment. Mice demonstrating 2 or more of pre-determined clinical signs were immediately culled. Surviving mice were culled at day 15 post infection.

#### Mouse sensitization

For sensitization experiments, BALB/c mice were utilised. Sensitized mice were subjected to 5 mosquitoes worth of saliva injections in 1μl of PBSA weekly for 4 consecutive weeks. Injections were made on dorsal side of left hind foot.

#### Mouse skin explants

Skin was dissected from the hind feet and transferred into 24 well tissue culture plate containing complete DMEM supplemented with 10% FCS, 10% TPB, 5 ml Pen/Strep and 5ml Glutamine broth. Explants were kept at 37°C with 5% CO_2_.

#### RNA purification and quantification

All tissue samples were lysed in 1ml Trizol reagent and shaken with 7mm stainless steel beads on a Tissue Lyser at 50Hz for 10min to ensure complete lysis of all tissues. 0.2ml chloroform was then added to all samples which were then inverted 15 times to allow for gentle mixing of the solutions. Afterwards, samples were centrifuged at 12,000g for 15min at 4°C in order to separate the mixture into a lower red phenol-chloroform phase and a colourless upper aqueous phase. The upper aqueous phase aqueous phase, containing the RNA, was transferred to a new tube containing an equal amount of 70% ethanol. RNA extractions were performed using the RNA mini purification kit (Life Technologies) by following the protocol provided with the kit. Purified RNA was then stored at −80°C.

Approximately 1μg of RNA in a volume of 9μl of RNAse free water was used for cDNA production using the “Applied Biosystems High Capacity RNA to cDNA” kit. Samples incubated at 37°C for 60min followed by heating to 95°C for 5min. The final cDNA was then stored at 4°C for short term use and at −20°C for long time storage.

Total cDNA was diluted 1 in 5, using RNAse free water and 1μl used per qPCR in 384 well plates. A master mix was made up of primers, water and SYBR green mix (perfecta, Quantabio.com). A triplicate technical replicate was made for each biological replicate. The generation of a standard curve was accomplished by the dilution of a 10^−2^ PCR-generated standard in a 10-fold serial dilution. A non-template control (NTC) consisting of RNAse free water and the master mix was also included. The PCR plates were run on the Applied Biosystems QuantStudio 7 flex machine.

Ct value was calculated automatically by the QuantStudio software which detects the logarithmic phase of the PCR reaction. Each samples relative quantity was calculated based on their position on the standard curve. The standard curve had to have an efficiency close to 100%, which was indicated by the coefficient R^2^≥0.998 and a slope of 3.3. Melt curves were conducted to control for primer specificity.

#### Plaque assays

BHK-21 cells in 12-well plates were grown to an 80% confluency and infected with virus serial dilutions prepared in 0.75% PBSA (PBS with 0.75% bovine serum albumin). 200μl virus dilutions was added to each well and left for an hour whilst rocking occasionally. 2xMEM medium supplemented with 4% FCS, 200 units/ml penicillin and 0.2 mg/ml streptomycin mixed 1:1 with 1.2% Avicel (FMC Biopolymer, UK), was added to the cells. Cells were incubated for 2 days at 37°C with 5% CO_2_. Cells were fixed in 10% PFA for an hour and stained with 0.1% Toluidine Blue for at least 30min. Plaques were counted and PFU was calculated per ml.

#### ELISA

ELISA was conducted using Mabtechs ELISA development kit. Plates were read on the Multiskan EX microplate reader (Thermo scientific) set to 450 nm to measure optical density (OD). Measurement was also taken at 540 nm and values were subtracted from 450nm measurements in order to correct for possible optical imperfections in the plate.

#### *In vitro* transepithelial electrical resistance (TEER) assay

Saliva was filter sterilised prior to use in endothelial cell culture. Human umbilical vein endothelial cells (HUVEC, Promocell) were cultured in Endothelial Growth Media (EGM-2, Promocell) at 37C in controlled atmosphere containing 5% CO_2_, until the culture reached 80% confluence. Cells were then detached by trypsinisation and 50,000 cells were seeded on each fibronectin-coated (final concentration 5μg/ml in PBS; Merck) Millicell Hanging Cell Culture Insert (PET membrane, pore size 0.4 μm, Merck Millipore) adapted for 24 well plates. Inserts were cultured in EGM-2 and TEER was measured every day using Millicell® ERS (Merck Millipore) until a plateau was reached, indicating the formation of a complete cell monolayer (normally 3-5 days after plating). 50% of the media was replaced with fresh EGM-2 every 2 days. Fresh medium containing Sialokinin (1μM) or mosquito saliva (0.5eq/ml; 1μl/ml) was added to both the upper and the lower chamber, and changes in TEER were monitored every 30 minutes for the first 3 hours, then every hour for a total of 8 hours. Before each measurement, cells were allowed to reach ambient temperature. Results were normalised against the values of untreated endothelial cells at each time point.

#### Tachykinin peptide searches

Searching the InterPro 82.0 database (October 2020) identified matches to the Tachykinin/Neurokinin-like, conserved site (PROSITE: PS00267, F-[IVFY]-G-[LM]-M-[G>]. InterPro: IPR013055) for 1’154 proteins from 219 deuterostomes but only 11 proteins from 7 protostomes. As well as *Ae. aegypti* Sialokinin, these included a peptide from a fungus-growing ant (Trachymyrmex cornetzi), four peptides from the extremely venomous Brazilian wandering spider (Phoneutria nigriventer), four peptides from three species of octopus (Eledoisins from Eledone cirrhosa, Eledone moschata, and Tachykinins 1 and 2 from Octopus vulgaris), and a likely false positive match to a protein from a tapeworm (Rodentolepis nana).

#### Sialokinin gene homology searches

Mosquito-focused VectorBase Release 48 (Giraldo-Calderon, et al. 2015) and insect-wide OrthoDB v10 (Kriventseva et al. 2019) gene family resources were searched with the prosialokinin precursor protein and corresponding gene (AAEL000229). VectorBase contains annotated genomes of 25 mosquitoes including *Ae. aegypti*, OrthoDB covers 148 insect species with 56 dipterans of which 17 are mosquitoes including *Ae. aegypti*, both resources also include two other culicine species: Ae. albopictus and *Culex quinquefasciatus*. Neither resource identified any homologues of AAEL000229 in any of the compared species. Protein sequence searches (BLASTp) of the NCBI non-redundant database returned only AAEL000229 itself (XP_001660125.1), and two other *Ae. aegypti* variants (AAD17916.1 and AAD16885.1). Protein sequence searches (tBLASTn) of all nucleotide data at VectorBase (genomes, transcriptomes, cDNAs, ESTs) identified AAEL000229 itself in the EST/cDNA and transcript databases, and in the *Ae. aegypti* AaegyL5 genome (Exon1: 5e-04 87%, Exon2 5e-04 100%, Exon3: 1e-29 87.7%) and the genome of the *Ae. aegypti* Aag2 cell line (Exon1: 5e-03 78.3%, Exon2: 5e-03 90%, Exon3: 3e-27 96.4%). No other credible hits were identified, the only hit with a comparable e-value (8e-04) was likely spurious as it was on the opposite strand of part of the coding region of a much longer gene encoding a DNA polymerase in *An. atroparvus* (AATE013717). The VectorBase region comparison tool, which uses genome-to-genome alignments to identify homologous and orthologous genomic regions between pairs of assemblies, identified no alignable regions in the *Ae. albopictus*, *Culex quinquefasciatus*, or *An. gambiae* genomes. We also searched (tBLASTn) the newly available assembly for Culex tarsalis (Main et al. 2020), which returned no significant hits.

#### dsRNA synthesis and injection into mosquitoes

A 173 bp fragment of the sialokinin coding DNA sequence (AAEL000229, AAEgL5) was amplified from *Ae. aegypti* Liverpool strain cDNAs with KOD Hot Start Master Mix (EMD Millipore) and sialokinin-specific primers with T7 RNA polymerase promoter sequence. The PCR product was purified using the QIAquick Gel Extraction kit (Qiagen). After sequencing, the PCR product was used as a template for a second PCR using the same primers and polymerase. For production of dsLacZ (used as control dsRNA), specific primers with T7 RNA polymerase promoter sequences were used to amplify a lacZ-derived fragment from plasmid template Drosophilaact5C-**β**Gal (Stock number 1220 obtained from DGRC) containing the E. coli lacZ gene. dsRNAs were synthesised and purified using the MEGAscript RNAi kit (Thermo Fisher Scientific) according to the manufacturer’s instructions. dsRNA was then purified and concentrated to 10 μg/μl in nuclease free water using 3M Sodium Acetate (Ambion) and ethanol precipitation. At 1 to 2 days after emergence, cold-anesthetised female mosquitoes were injected into their thorax using a nanoinjector (Nanoject II, Drummond Scientific) with 2 μg of dsRNA (dsSialokinin or dsLacZ). Four days post-injection, saliva was collected from dsLacZ-(control) and dsSialokinin-injected females (pools of saliva from 5 females) and after salivation, females (pooled accordingly to saliva pools) were sampled in 1ml of TRIzol (Invitrogen) and stored at −80°C until RNA extraction.

### Mosquito RNA extraction and RT-qPCR

Females were homogenized (Precellys 24, Bertin Technologies) in TRIzol (Invitrogen) and samples were centrifuged at 6500g for 30 sec. Total RNA was extracted using the TRIzol method according to the manufacturer’s (Invitrogen) protocol except that 1M 1-Bromo-3-ChloroPropane (BCP) (Sigma-Aldrich) was used instead of chloroform. DNase treatment was performed during 30 min at 37°C following the manufacturer’s protocol (TURBO DNase, kit Invitrogen), except that RNasine 0.36 U/μL (Promega) was also added. cDNAs were synthesized using total RNA (25 ng/μL) and M-MLV Reverse Transcriptase (Invitrogen). All cDNAs were aliquoted and stored at −20°C until qPCR. qPCR assays were performed with the Fast SYBR Green Master Mix method (Thermo Fisher Scientific) according to the manufacturer’s protocol and using specific primers (Sigma-Aldrich) for genes of interest; reactions were run on an ABI 7500 Fast RT PCR machine and results were analysed with the 7500 Software v2.0.6. To quantify sialokinin knockdown efficiency, data were analysed using the comparative Ct (cycle threshold) method using S7 ribosomal protein gene as a standard gene for normalisation. One of the dsLacZ sample was set to RQ=1 and all other samples expressed relatively to this sample.

#### Sialokinin peptide generation

Sialokinin peptide (N-T-G-D-K-F-Y-G-L-M-amide) was synthesised de novo to >95% purity by Cambridge Research Biochemicals

#### Statistical analysis

Analysis of RT-qPCR data was done with Microsoft Excel by the use of the median of the technical replicates and normalizing them to the median of the technical replicates of the housekeeping genes. All data was analysed with GraphPad Prism software. Non-parametric Kruskal-Wallis test was used for comparisons between more than two groups whilst non-parametric Mann-Whitney was used for comparisons between two groups. Ordinary-ANOVA was performed for comparisons between more than two groups of normally distributed data. Analysis of survival curves was conducted using the logrank (Mantel Cox) test. All differences were considered significant at p<0.05. All plots have statistical significance indicated as follows: *p<0.05, **p<0.01, ***p<0.001, ****p<0.0001, ns=not significant.

---

**sFigure 1.**
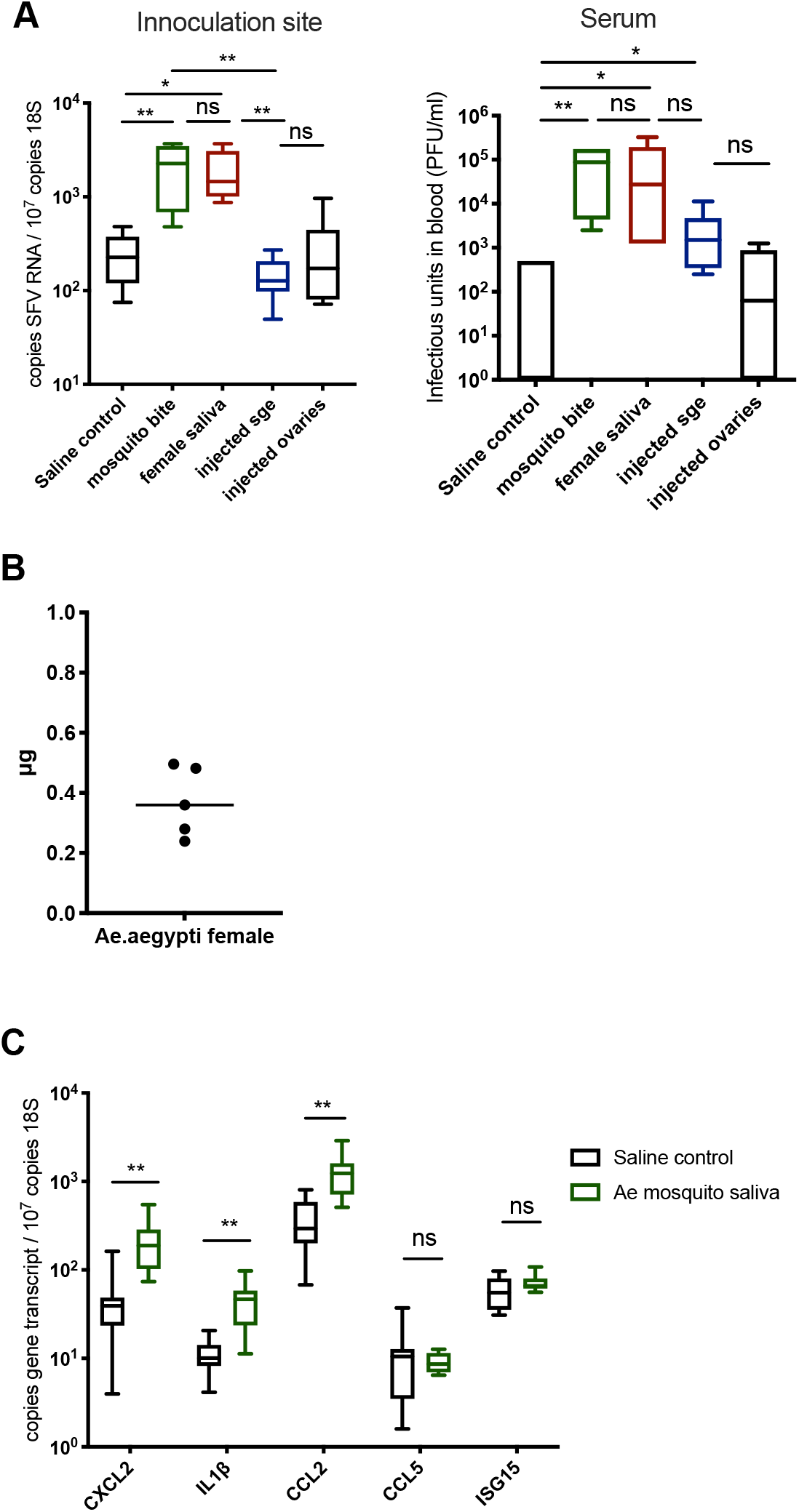
(A) Mouse skin was inoculated with 10^4^ PFU of SFV4 alone or alongside 1.86μg saliva, 5 denatured salivary glands or ovaries, or exposed to up to 5 bites from Ae.aegypti. (B) Saliva acquired from 5 mosquitoes were pooled and protein content was quantified via nanodrop. Each dot represents the average protein concentration per mosquito. (C) Mouse skin was injected with either saline control or 1.86μg saliva of Ae.aegypti. Copy number of host transcripts in the skin was determined by qPCR at 6 hours (n=6)

**sFigure 2.**
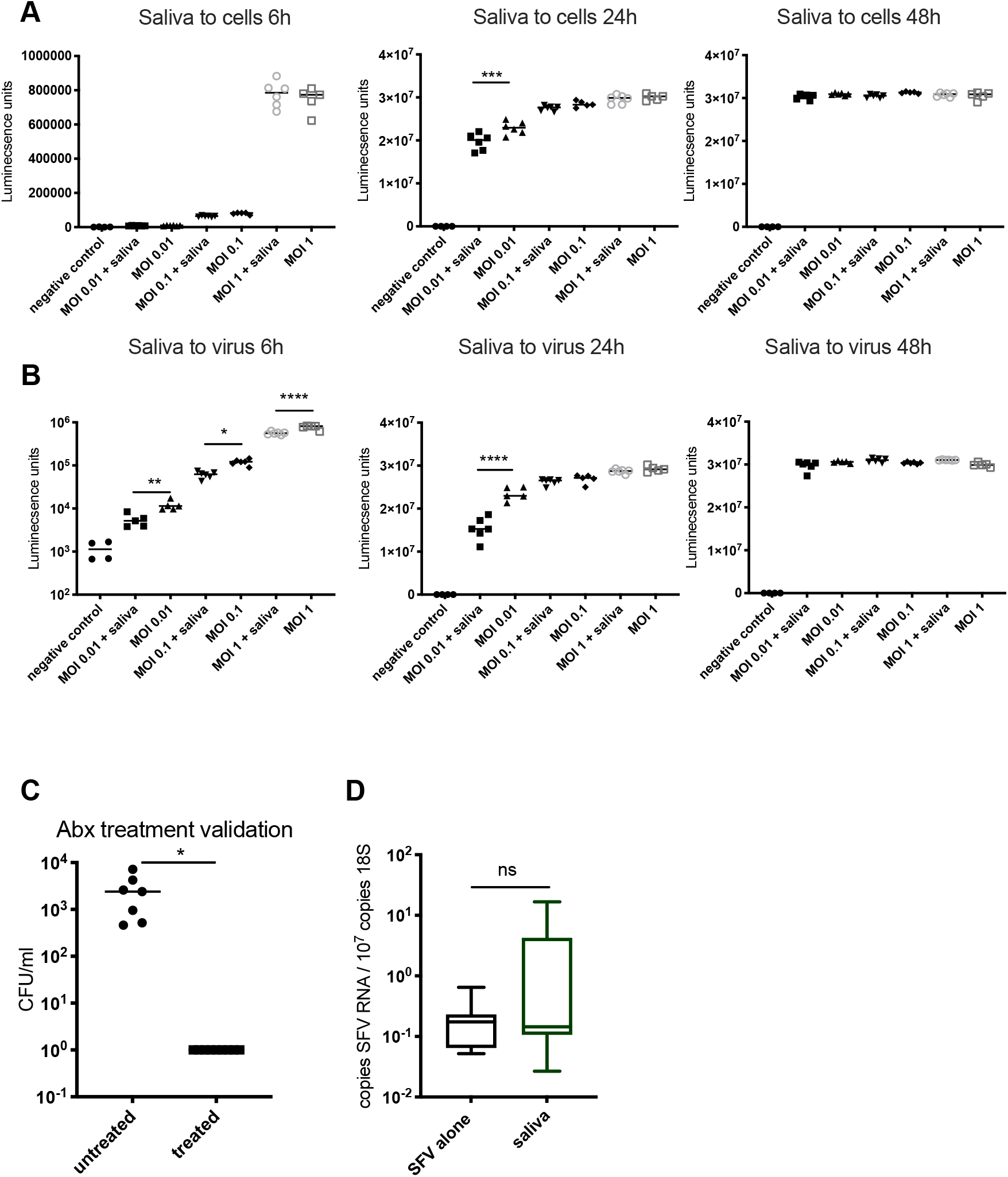
(A-B) Primary cultures of dermal fibroblasts (target of SFV in vivo) where infected with Gluc-expressing SFV at MOIs of 0.01, 0.1 and 1 and luciferase activity of tissue culture supernatant assayed at 6, 24 and 48 hpi. (A) Cells were pre-treated with saliva for 1h prior to infection (n=6). (B) Saliva was added to virus prior to infection of cells (n=6). (C) Efficacy validation of antibiotic treatment from pupae stage onwards (pen/strep at 200 U per ml, gentamycin at 200 μg/ml, and tetracycline at 100 μg/mL). Mosquitoes at 2 weeks post emergence were dipped in ethanol, dried and mosquito extract plated on agar plates in 10-fold dilutions. CFU/ml was calculated at 24h. (D) Mouse skin was bitten by Aedes mosquitoes and inflammation allowed to develop for 16 hours, then biopsies take and infected ex vivo with 10^4^ PFU SFV. Viral RNA and host 18S were quantified by qPCR (n=8).

**sFigure 3.**
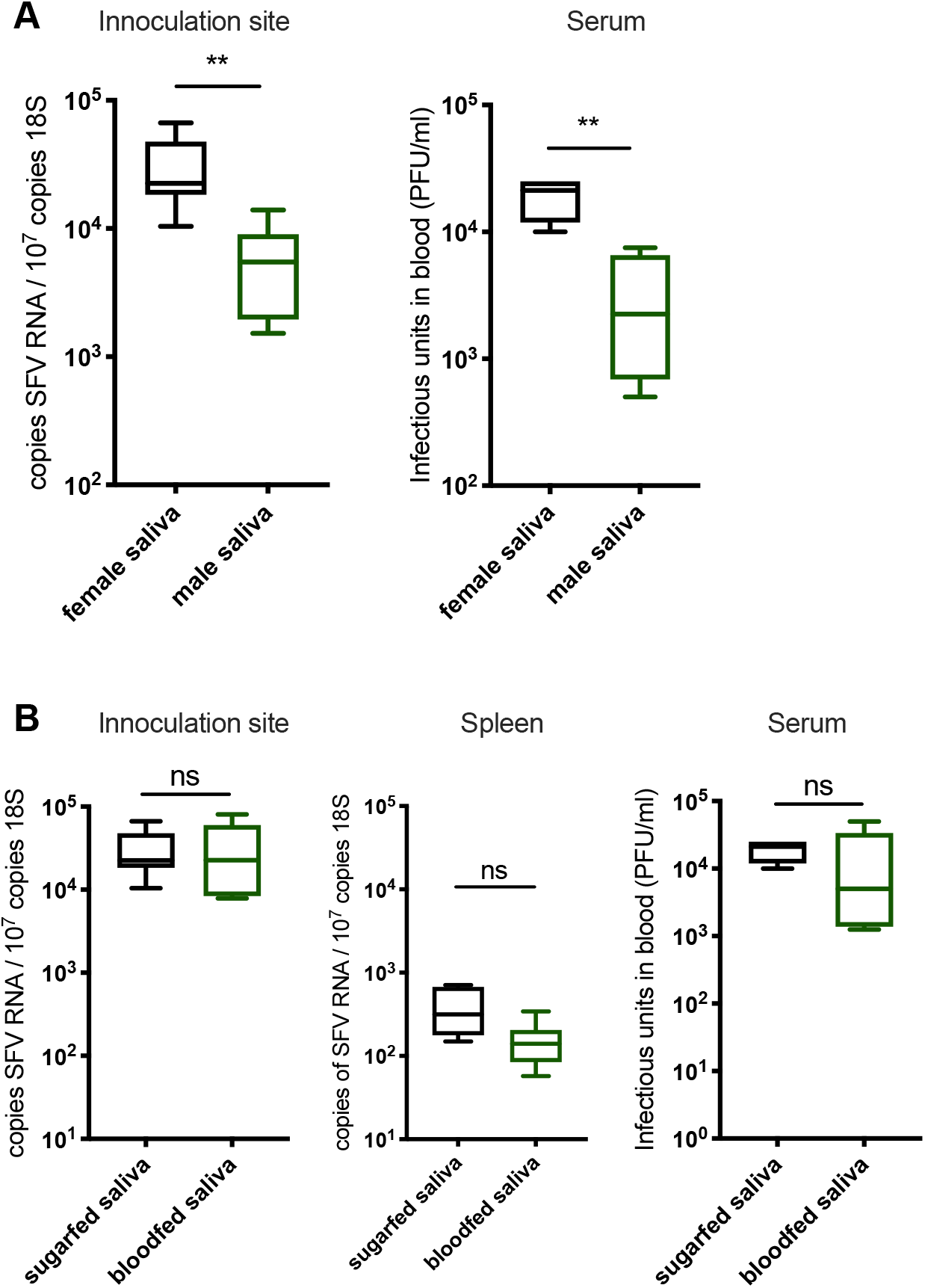
(A,B) Mouse skin was inoculated with 10^4^ PFU of SFV4 alone or with Ae.aegypti saliva in the upper skin of the left foot. Viral RNA and host 18S were quantified from skin and spleen by qPCR and viral titres of serum by plaque assays at 24hpi. (A) 1 μl male or female Ae.aegypti saliva in PBSA, derived from mosquitoes reared in the same cage. Because male saliva contained less total protein, protein content was normalised by diluting female saliva with PBSA prior to injection (n=6). (B) Saliva from bloodfed or exclusively sugarfed female Ae.aegypti mosquitoes. (n=6)

**sFigure 4.**
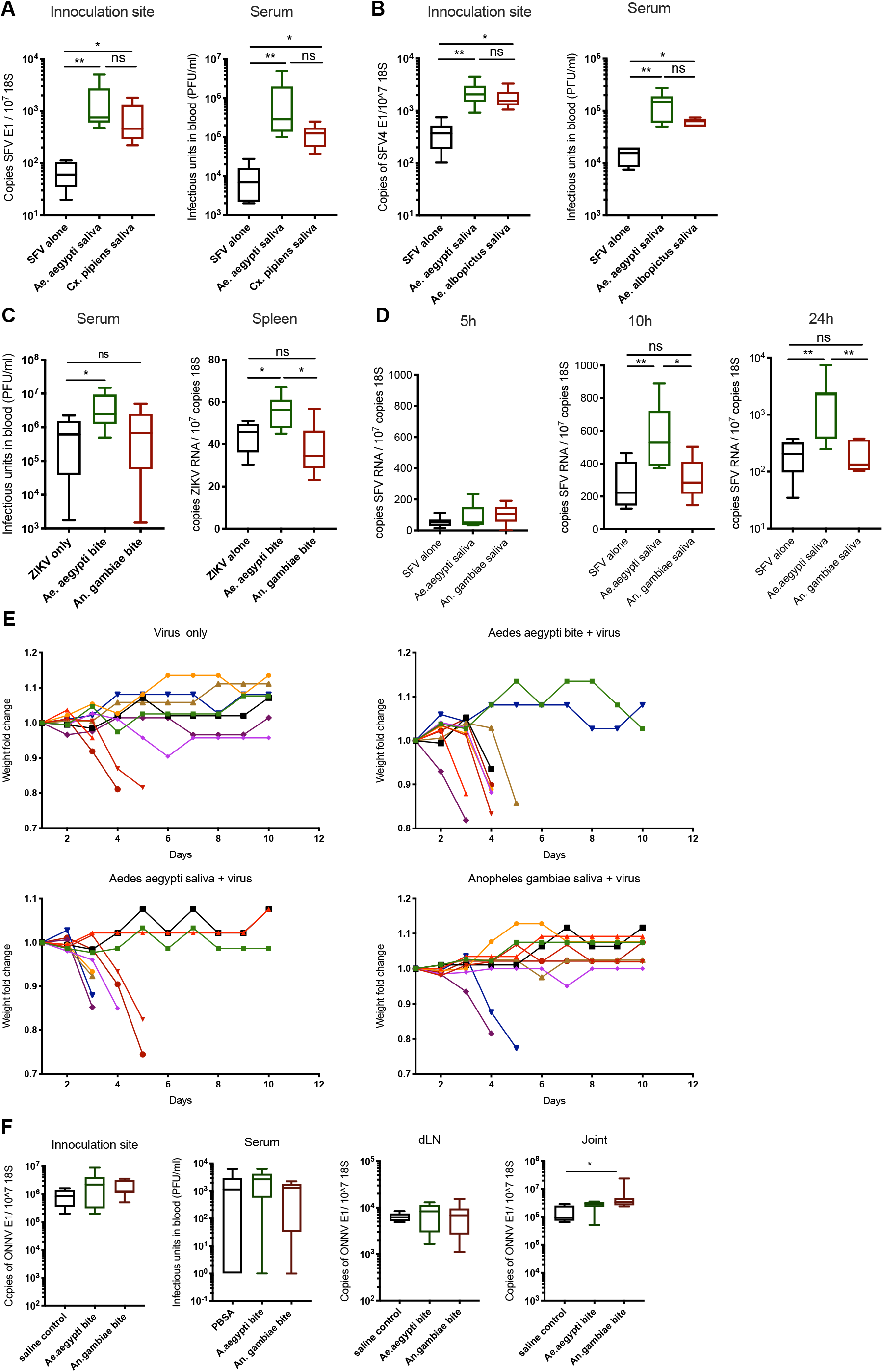
(A-E) Mouse skin was inoculated with 10^4^ PFU of SFV4 alone or alongside 1.86μg saliva of either Ae.aegypti, Ae.albopictus, Cu.pipiens or An.gambiae. SFV RNA and host 18S and serum viral titres were quantified at 24 hpi. (C) Mouse skin was exposed to up to 3 bites of either Ae.aegypti or An.gambiae mosquitoes. (D) SFV RNA and host 18S and serum viral titres were quantified at 5, 10 and 24 hpi. (E) Weights of mice from survival experiment (Fig 4B). Mice were weighed once daily. (F) Mouse skin was infected with 2 ×10^5^ PFU ONNV alone or alongside 1.86μg saliva or following up to 3 bites of either Ae.aegypti or An.gambiae in the upper skin of the left foot. ONNV RNA and host 18S from tissues were quantified by qPCR and serum viral titres were quantified via plaque assays at 48 hpi.

**sFigure 5.**
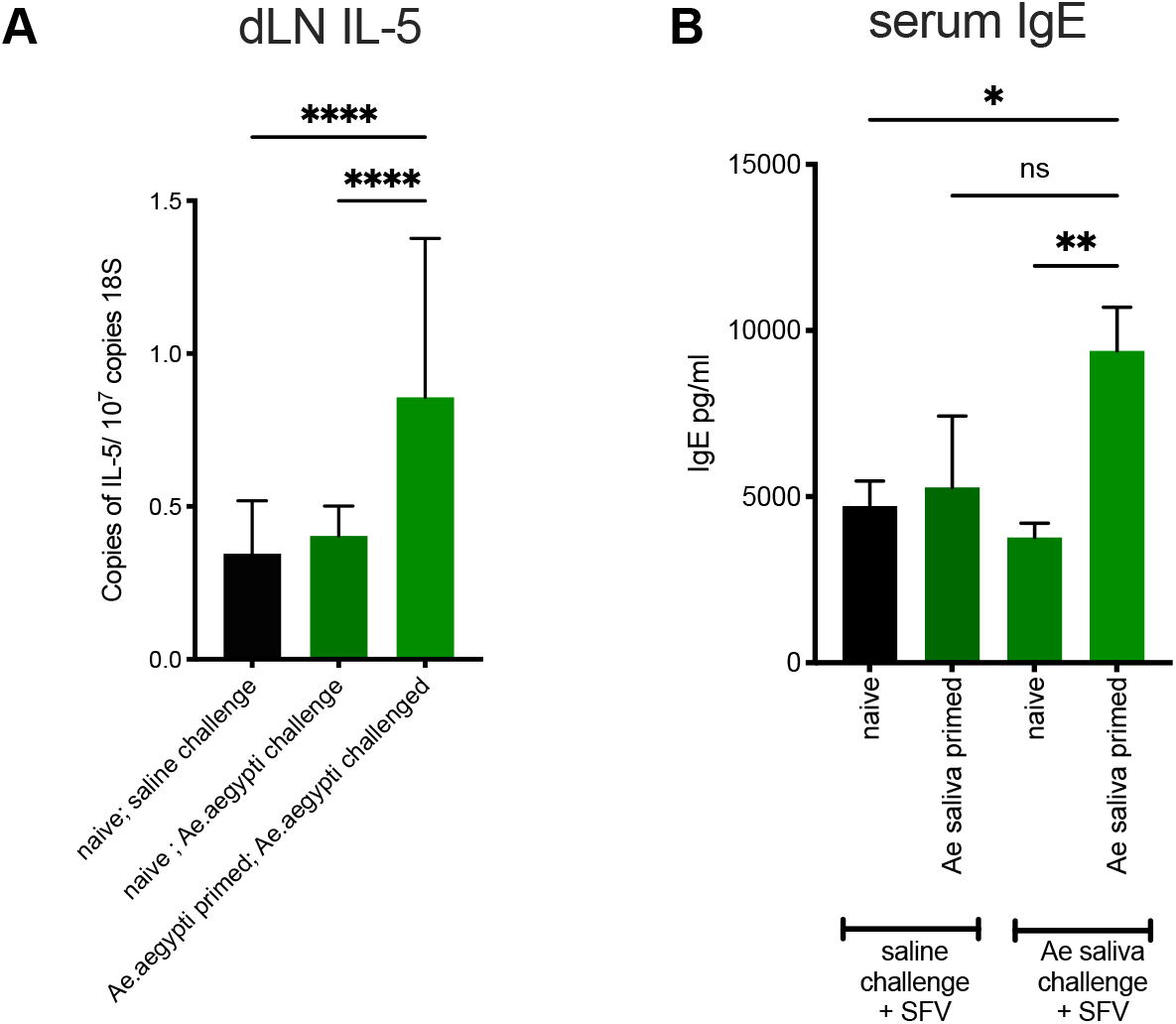
(A,B) Balb/c mice skin was inoculated with 10,000 PFU of SFV alone or with Ae. aegypti saliva. Mice were either naïve to saliva or primed to saliva by injections of mosquito saliva weekly for 4 consecutive weeks. (A) Draining popliteal lymph node IL-5 transcript expression at 2 hpi were quantified by qPCR (n>6). (B) Serum total IgE was quantified at 2 hpi by ELISA (n>6).

**sFigure 6.**
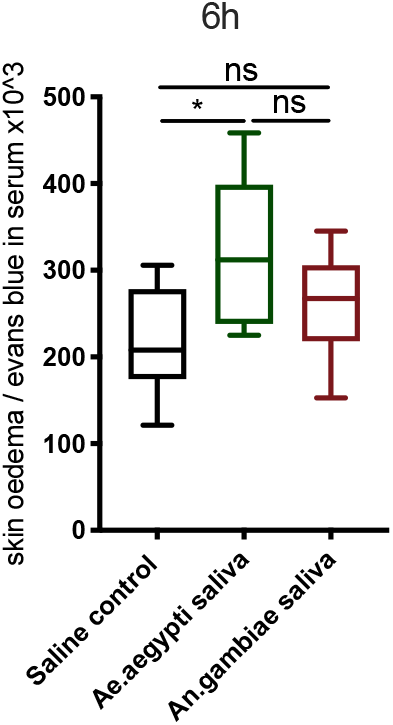
Mice administered i.p. with Evans blue, were injected with 1.86μg of mosquito saliva in the skin of naïve or sensitised mice of either Ae.aegypti or An.gambiae. Extent of oedema assessed by quantification of Evan’s blue dye leakage into skin at 6h post saliva via colorimetric assay (n=6).

**sFigure 7.**
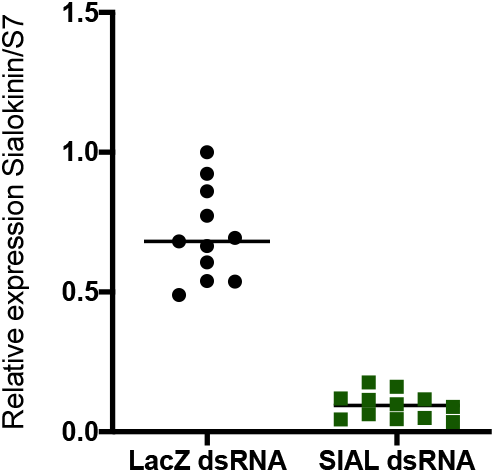
Knockdown efficiency of sialokinin expression in females Ae. aegypti. Expression levels of sialokinin in females previously injected with dsRNA targeting lacZ or sialokinin. Data were analysed using the comparative cycle threshold method using S7 ribosomal protein gene as a standard gene for normalisation. One of the dsLacZ sample was set to RQ=1 and all other samples expressed relatively to this sample. Median plus interquatile range shown. The expression of sialokinin was efficiently knockdown (87% median reduction) in dsSialokinin-injected females compared to dslacZ- ones (Mann-Whitney test, p value <0.0001, N = 11 and 12 pools of 5 females for dsLacZ and dsSialokinin respectively).

